# A theoretical framework for multi-species range expansion in spatially heterogeneous landscapes

**DOI:** 10.1101/2021.11.09.467881

**Authors:** Lukas Eigentler, Nicola R. Stanley-Wall, Fordyce A. Davidson

## Abstract

Range expansion is the spatial spread of a population into previously unoccupied regions. Understanding range expansion is important for the study and successful manipulation and management of ecosystems, with applications ranging from controlling bacterial biofilm formation in industrial and medical environments to large scale conservation programmes for species undergoing climate-change induced habitat disruption. During range expansion, species typically encounter competitors. Moreover, the environment into which expansion takes place is almost always heterogeneous when considered at the scale of the individual. Despite the ubiquitous nature of these features, the impact of competition and spatial landscape heterogeneities on range expansion remains understudied. In this paper we present a theoretical framework comprising two competing generic species undergoing range expansion and use it to investigate the impact of spatial landscape heterogeneities on range expansion with a particular focus on its effect on competition dynamics. We reveal that the area covered by range expansion during a fixed time interval is highly variable due to the fixed landscape heterogeneities. Moreover, we report significant variability in competitive outcome (relative abundance of a focal species) but determine that this is induced by low initial population densities, independent of landscape heterogeneities. We further show that both area covered by range expansion and competitive outcome can be accurately predicted by a Voronoi tessellation with respect to an appropriate metric, which only requires information on the spatial landscape and the response of each species to that landscape. Finally, we reveal that if species interact antagonistically during range expansion, the dominant mode of competition depends on the initial population density. Antagonistic actions determine competitive outcome if the initial population density is high, but competition for space is the dominant mode of competition if the initial population density is low.

## 1 Introduction

The spread of a population into space previously unoccupied by that population is commonly referred to as *range expansion*. This is a ubiquitous phenomenon; for example, range expansion occurs during growth of microbial colonies (Buttery et al. 2012; Hallatschek et al. 2007), ecological invasions (Fraser et al. 2015; Hastings et al. 2005; Okubo et al. 1989; Pejchar and Mooney 2009), the spread of epidemics (Artois et al. 2018; Diekmann 1979) and even human migration (Moreau et al. 2011; Templeton 2002). Notably, range expansion is not just a feature of the early history of a species (e.g. spread of a new disease), but also occurs due to changes to the environment such as those induced by climate change (with habitats typically shifting towards poles or higher elevations) (Hill et al. 2001; Rosenzweig et al. 2007; Vos et al. 2008; Wilson et al. 2009). Under current policies, climate change is predicted to accelerate species extinction and is estimated to be currently threatening over 15% of species globally (Urban 2015). Therefore, better understanding how range expansion enables species to track shifts of habitable environments is becoming increasingly important (Rosenzweig et al. 2007). Understanding the fundamental dynamics that govern range expansion will provide insights into the resilience of species to climate change (MacDonald and Lutscher 2018) and could provide opportunities for landscape management as part of conservation programmes (Vos et al. 2008; Wilson et al. 2009). As range expansion typically takes place over large spatial scales (compared to the size of an individual), populations are likely to encounter spatially heterogeneous landscapes (Crone et al. 2019; Fraser et al. 2015; Hill et al. 2001; With 1997; With 2002).

Most ecological systems are underpinned by competitive dynamics between species (Levine and HilleRisLambers 2010; Valladares et al. 2015). Therefore, during range expansion, populations typically comprise different species that compete for space and resources by means of spatial expansion and other competitive interactions. Previous studies have addressed many different facets involved in multispecies range expansion; for example, intraspecific competition has been shown to slow the speed of range expansion and also affect the shape of the population fronts (Legault et al. 2020); successive range expansion of different species has been shown to enable coexistence with a fractal-like structure in the population (Goldschmidt et al. 2017); classical results on non-transitive competitive hierarchies were highlighted to not necessarily apply to multi-species range expansion (Weber et al. 2014); and genetic drift at population fronts during range expansion has been shown to lead to a loss of genetic diversity (Hallatschek et al. 2007). While the role of spatial heterogeneity in the landscape on evolutionary dynamics in range expansion has previously been investigated (Gralka and Hallatschek 2019; Wegmann et al. 2006), its impact on competition for space and its interplay with other interspecific competition dynamics during range expansion remain understudied.

From a theoretical perspective, competition dynamics in biological and ecological systems are most commonly characterised by their asymptotic behaviour - often an equilibrium state (e.g. Chesson (2000) and McPeek (2012)). However, such an approach is not suitable for description of range expansion as this asymptotic approach fails to account for the importance of the invasion dynamics (Ghosh et al. 2015; Travis and Dytham 2002). Instead, non-equilibrium analyses of range expansion have been employed that are restricted to a finite time interval with results typically reported from a fixed, predefined endpoint (Goldschmidt et al. 2017; Hallatschek et al. 2007; Legault et al. 2020; Weber et al. 2014). The importance of space to competition dynamics in equilibrium settings is a well-explored topic. Spatially-extended dynamics are known to enable species coexistence at equilibrium in some cases in which competition in well-mixed conditions would lead to competitive exclusion. One classical example is a trade-off between dispersal abilities and local competitiveness (Horn and MacArthur 1972; Levins and Culver 1971). Such a trade-off creates behavioural niches (e.g. Whittaker et al. (1973)), which enables a locally weaker species to persist in a population if it is able to colonise new areas more rapidly than its locally superior competitor(s) (Tilman 1994). Thus, a trade-off between local competitiveness and colonisation abilities creates a balance that enables coexistence through spatial segregation (Gravel et al. 2010; Hassell et al. 1994; Horn and MacArthur 1972; Levins and Culver 1971; Tilman 1994). Similarly, coexistence in a spatially segregated equilibrium state can also be induced by non-transitive competitive hierarchies of three or more species, usually referred to as “rock-paper-scissor dynamics” (e.g. Avelino et al. (2019), Kerr et al. (2002), Lowery and Ursell (2019) and Reichenbach et al. (2007)).

Many antagonistic competitive interactions that take place in a spatially extended context require, at a fundamental level, spatial co-location. Hence, competition for space and competition through antagonistic actions are intrinsically linked. To understand the precise dynamics of competitive mechanisms affecting range expansion, it is therefore key to first attain knowledge about the role of spatial dynamics, in particular those that lead to spatial segregation. In a previous paper, we investigated the role of competition for space between two biofilm-forming bacterial strains in the specific case of microbial range expansion on spatially homogeneous substrates (Eigentler et al. 2021). In brief, we revealed that the initial population density has a significant impact on competitive outcome (defined to be the relative abundance of one focal species across the whole community). Starting with a fixed 1:1 ratio between populations, high initial densities consistently led to an equal competitive outcome. By contrast, low initial population densities resulted in highly variable competitive outcomes. Furthermore, in (Eigentler et al. 2021) we defined a predictive tool that could be used to determine competitive outcome based solely on the distribution of the initial population. In short, the method is as follows. A circle was first drawn around the initial population. Next, for each point on the circle, the closest initial population patch was determined and the point on the circle labelled correspondingly. Finally, this labelling was used to define an index, termed the *access to free space score*, that quantified the proportion of points on the circle associated with each strain (Fig. 1.1). For fixed initial population density, we revealed a remarkably strong linear correlation between the access to free space score and the competitive outcome of a strain.

**Figure 1.1:**
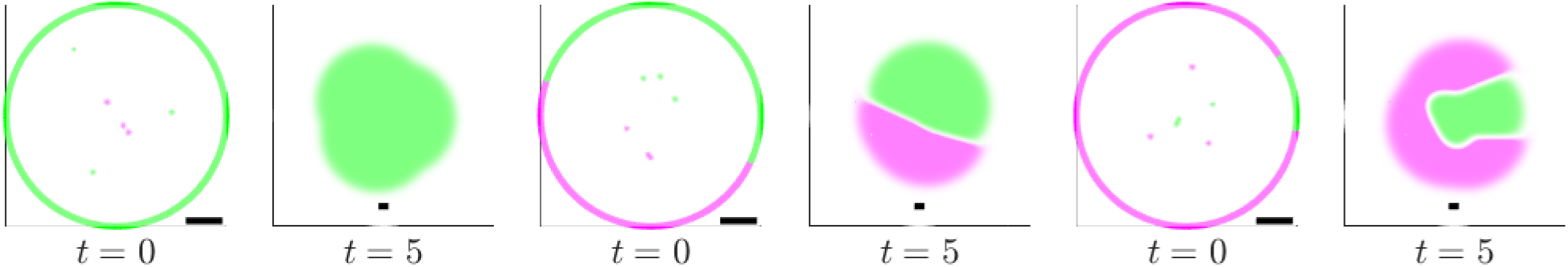
Predictions of competitive outcome in a spatially homogeneous landscape. Three example realisations of (2.1) on a spatially homogeneous landscape with *N* = 6 initial population patches are shown. The parameter values are *r*_1_ = *r*_2_ = 5, *d*_1_ = *d*_2_ = 0.1, *c*_21_ = 10 and *c*_12_ = 50. Note that these parameter values mean that both species are governed by the same growth dynamics in the absence of a competitor, but that *B*_2_ (green) is the intrinsically stronger species under competition. The initial condition is classified by drawing a circle around the initial population and determining the closest initial patch to each point on the circle (see text). Note that the visualisation of the initial condition shows a blow-up of the domain centre only. The scale bar is one unit length in all figures. This figure is adapted from (Eigentler et al. 2021) in accordance to its CC-BY-NC-ND 4.0 International license.

**Figure 2.1:**
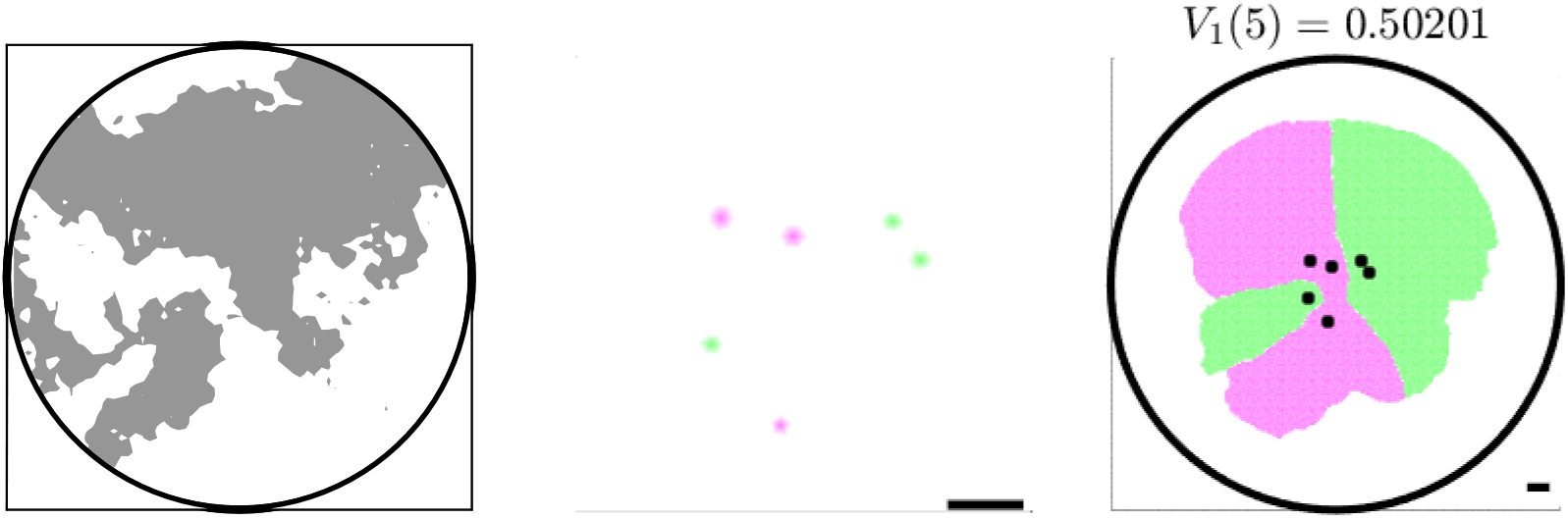
Voronoi tessellation with respect to front propagation metric. A Voronoi tessellation, i.e. the sets 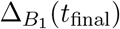 and 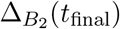, where *t*_final_ = 5, with respect to the front propagation metric *t*_FP_ is shown (right). The initial cell patches are shown by black markers and are also visualised separately (middle). Note that the middle column shows a blow-up of the centre of the computational domain only. The scale bars are one unit length long. The underlying spatially heterogeneous domain is shown in the left column. Grey areas indicate good environmental conditions with *r* = 5, *d* = 0.1 and white areas indicate challenging environmental conditions with *r* = 2.5, *d* = 0.05.

In this study, we focus on the interplay of competition for space and antagonistic interactions in multi-species range expansion that takes place in spatially heterogeneous landscapes. We first confirm that, all else being equal, changes in a spatially heterogeneous landscape across different realisations of range expansion represent an additional source of variability. Based on this observation, we investigate (a) whether the initial population density remains the determinant of variability in competitive outcome or whether spatial heterogeneities become dominant; (b) if predictions of spatial spread and competitive outcome during range range expansion can be made based on the initial population distribution despite the heterogeneities in the landscape; and (c) whether competition for space or antagonistic interactions are the dominant competitive mode during range expansion or whether there is a fundamental dependence on the spatially heterogeneous landscape.

We first present the mathematical framework and methods used in the model analysis in Section 2. The results of our model analysis are presented next in Section 3, where we first focus on competition for space (Section 3.1). Then, we investigate the impact of the interplay of spatial dynamics and antagonistic mechanisms (Section 3.2). Finally, we discuss the implications of our results in Section 4.

## 2 Theoretical framework

### 2.1 Model

Multi-species range expansion can be abstracted to the interplay between the key processes of (net) local growth, dispersal and interspecific interactions. We capture these dynamics in a mathematical framework and account for spatial heterogeneity in the landscape by employing space-dependent dispersal coefficients and growth rates. These heterogeneities represent variations in environmental conditions, such as the availability of growth limiting nutrients.

The model describes the dynamics of two generic species *B*_1_(**x**, *t*) and *B*_2_(**x**, *t*) and tracks their dynamics using as system of partial differential equations (PDEs) based on the spatially extended competitive Lotka-Volterra equations:

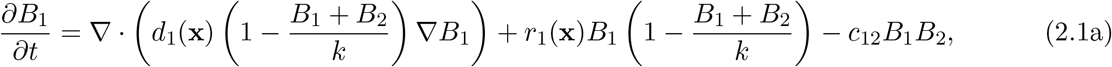

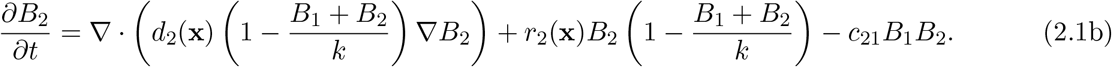

Here **x** ∈ Ω is a point in a two-dimensional circular spatial domain Ω = {**x** ∈ ℝ^2^ : ∥*x*∥*< R*_Ω_} for some positive constant *R*_Ω_ and time *t* > 0. To preserve generality, we choose a simple, logistic law as a representative form of local population growth, where *r*_1_(**x**) ≥ 0 and *r*_2_(**x**) ≥ 0 denote the maximum growth rates of *B*_1_ and *B*_2_, respectively, and *k* ≥ 0 the carrying capacity. Note that growth is limited by the total population density and again for simplicity we assume the carrying capacity to be constant (see Discussion). This represents growth limitation due to intraspecific and interspecific competition for resources, such as nutrients or space. Dispersal is assumed to be random and is modelled by a diffusion term. However, diffusion of either species into occupied territories is assumed to be limited by the resident population. This limitation in dispersal appears to be a common feature of interacting systems across a range of scales. Examples include interacting microbial populations (Matoz-Fernandez et al. 2020; Stefanic et al. 2015) and territorial animal species (Cozzi et al. 2018). Limitation is accounted for in the model through density-dependent flux terms that decrease with total population density from their respective maxima *d*_1_(**x**) ≥ 0 and *d*_2_(**x**) ≥ 0 at *B*_1_ + *B*_2_ = 0 to zero at *B*_1_ +*B*_2_ = *k*. Moreover, upon contact, species are assumed to engage in antagonistic actions leading to additional death or growth reduction between both species not accounted for by the logistic growth terms. The constants *c*_12_ ≥ 0 and *c*_21_ ≥ 0 denote the respective antagonistic rates. For brevity, we refer to these antagonistic interactions as “killing terms”. However, we note that that these interaction terms could be assimilated into the standard form of the Lotka-Volterra equations with intraspecific competition coefficients *r*_1_(**x**)/*k* and *r*_2_(**x**)/*k* and interspecific competition coefficients *r*_1_(**x**)/*k + c*_12_ and *r*_2_(**x**)/*k + c*_21_ in (2.1a) and (2.1b), respectively. Thus, the model makes no explicit assumption on mode of the interaction between the species with the exception that the impact of interspecific competition is assumed to be stronger than or equal to the impact of intraspecific competition for each species.

### 2.2 Model initial conditions and simulation

The model (2.1) is solved using a finite element method employed by the `PDE Toolbox` in Matlab (The MathWorks Inc. 2020). Numerical integration is stopped before the population reaches the boundary of the computational domain Ω. In this way, neither the explicit form of the boundary conditions nor the shape of the boundary bear influence on the model solution. For computational ease, we assume Ω to be a disc and set ∇*B*_1_ · **n** = ∇*B*_2_ · **n** = 0, where **n** denotes the outward normal on *ᑯ*Ω.

The initial population is assumed to be confined to a subset 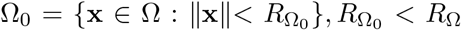 in the centre of the computational domain Ω. High initial population densities are represented by spatially homogeneous initial conditions in Ω_0_, i.e. *B*_1_(**x**, 0) ≡ *k*_1_, *B*_2_(**x**, 0) ≡ *k*_2_ with *k*_1_ + *k*_2_ ≤ *k*. The representation of very low initial population densities requires a different approach; at such densities, individuals are not necessarily spread uniformly in space. For example, inoculation of low densities of bacterial cells initially leads to the formation of small, spatially segregated microcolonies (Eigentler et al. 2021), and invasion of mammal species into previously unoccupied habitats typically originate from a small number of individuals (Campbell et al. 2012; Middleton 1930). To represent this initial clustering in the model, each species is initially confined to a number of small patches within Ω_0_ in which their density is set to carrying capacity. Unless otherwise stated, the locations of these initial patches are chosen uniformly at random in Ω_0_ with no overlap and their total number is denoted by *N* ∈ ℕ. For brevity, we refer to this type of initial condition as a *patch initial condition* throughout the manuscript. Computationally, the spatial mesh used to numerically solve the model imposes restrictions on the location and size of such patches. Therefore, we randomly choose the initial patch positions for each species from the set of mesh nodes in Ω_0_ and set that species to carrying capacity at the nodes and to zero everywhere else. It is noted that depending on the spatial scale and precise application of the model, initial patches created by this method may be larger than the size of a single individual. In these cases, it is reasonable to assume that such patches can be identified with single individuals that were initially able to reproduce without interaction with competitors.

### 2.3 Competitive outcome

For species *B*_1_, we define *the competitive outcome* of multi-species range expansion to be a time-dependent function, 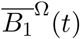, quantifying the relative abundance of species *B*_1_ across the whole computational domain at time *t* > 0 and given by

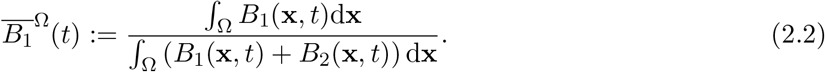

We focus mainly on competitive outcome at the chosen endpoint *t* = *t*_final_ of the model integration, but refer to the temporal dynamics of this quantity where appropriate. The chosen value of *t* = *t*_final_ is sufficiently large to ensure the area covered by range expansion is significantly larger than the area of Ω_0_ in which the initial population patches are placed, but sufficiently small so that the population does not reach the boundary of the computational domain Ω during the simulation. Note that competitive outcome for *B*_2_, denoted by 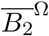, is given by 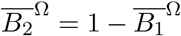. Due to this simple relationship and for ease of exposition, throughout the paper we only refer to the competitive outcome for species *B*_1_.

### 2.4 Area covered by range expansion

We further quantify the *area covered by range expansion*. Like competitive outcome, the area covered by range expansion, *A*(*t*), is a time-dynamic quantity, defined by

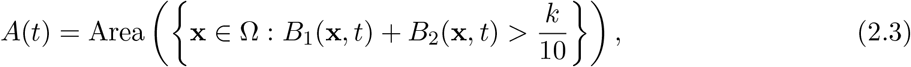

where Area(·) denotes the area of a set in the Euclidean sense.

### 2.5 Front propagation metric

The relation between Euclidean distance and propagation time used in defining the access to free space score in homogeneous environments does not hold in spatially heterogeneous landscapes. Therefore, in this section, we define new tools to predict competitive outcome that take into account spatial heterogeneities in the landscape.

In order to make predictions regarding competitive outcome, we first determine the times that individuals in a small patch located at **x** ∈ Ω would take to move along each possible path from **x** to some distant point **y** in the absence of any competitive interaction. We then define the *front propagation metric* to be the shortest such time taken. A rigorous definition is as follows. Denote by 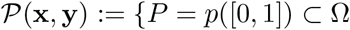, where 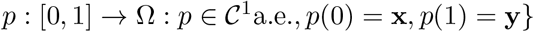 the set of all paths from **x** to **y**. For a given path 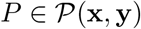, the time taken to move along the path is given by

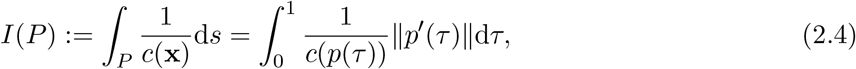

where 0 ≤ *c*(**x**) < ∞ represents the propagation speed along the path *P*. We define the *front propagation metric* from **x** to **y** as

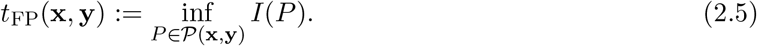

We show in the supplement (Section S2) that *t*_FP_(**x**, **y**) is a metric in the mathematical sense.

Note that for our definition of the front propagation metric *t*_FP_ to hold, certain conditions are required on *r*(**x**) and *d*(**x**) to guarantee that the set 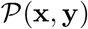 covers all relevant connections between points **x** and **y**. Choosing *r*(**x**) and *d*(**x**) to be differentiable would be sufficient and we believe these conditions could be considerably relaxed. Moreover, we note that for computational calculations of *t*_FP_ (more details below), spatial structure of the landscape is required to occur on the same (or coarser) scale than that of the domain discretisation to ensure accuracy of the computation. However, theoretically, the definition may fail on landscapes that have, for example, a full fractal structure to an infinitesimal level. In such cases, it is not clear that the infimum exists in (2.5). Hence and given that we are principally interested in the numerical implementation of the metric, we do not pursue a definitive set of necessary conditions on the parameters here.

It is well known that *pulled* travelling wave solutions (that is, wave fronts whose dynamics are governed only by their leading edges where linear terms dominate) exist for system (2.1) i<u>n</u> spatially homogeneous domains. The minimum speed of these travelling waves is given by 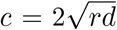. It can be shown that a large class of front solutions to system (2.1) asymptotically approach the dynamics of this minimum speed travelling wave (Stokes 1976). For spatially heter eneous landscapes, we therefore reasonably assume the *local* front propagation speed to be 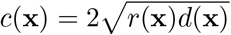 for all **x** ∈ Ω. [Numerical simulations confirm that this is indeed a reasonable approximation.] Finally, we assume that both species undergo the same growth and diffusion dynamics in the absence of a competitor species (i.e. *r*_1_(**x**) = *r*_2_(**x**) =: *r*(**x**)) and *d*_1_(**x**) = *d*_2_(**x**) =: *d*(**x**)). In this case, the expansion fronts of both species are governed by the same expansion speed. We provide more information on relaxing this assumption in the Discussion section and Section S4.

Numerically, *t*_FP_ is computed by discretising the computational domain Ω into a weighted graph and applying Dijkstra’s algorithm (see SI for details). Nodes of the graph are defined to be the centres of the finite elements of the domain discretisation. Graph edges connect graph nodes of elements that share edges of the finite elements. Graph edges are weighted by the time a front would require to travel along the edge, with its speed being approximated by the mean of the front speeds *c*(**x**) evaluated at both nodes. In other words, a graph edge connecting graph nodes at positions **x** and **y** is assigned weight 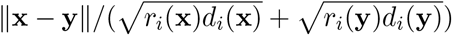.

### 2.6 Voronoi tessellations

We now define a classification of the initial condition by performing a Voronoi tessellation with respect to the metric *t*_FP_. That is, given *N* ∈ ℕ initially occupied patches centred at **x_1_**, … **x_N_** ∈ Ω_0_ we define the sets 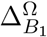 and 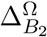 with 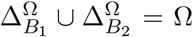 to comprise points closest to initial patches of *B*_1_ and initial patches of *B*_2_, respectively, namely,

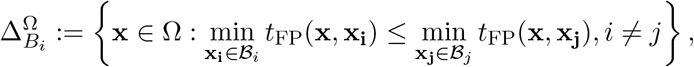

where 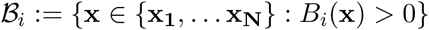, *i* = 1,2.

Next, we define the sets

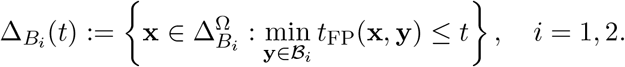

These sets provide an estimate for regions in Ω occupied by each species at any given time *t* (with the proviso that no competitive interactions take place).

Finally, we define the *Voronoi index V_i_*(*t*) by

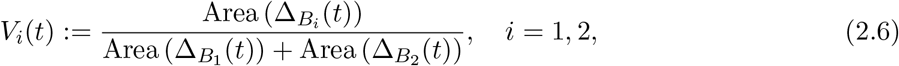

where Area(·) denotes the total area of a set (or collection of sets) in the Euclidean sense. The Voronoi index allows the classification of the initial condition for species *B*_*i*_ using a single number *V*_*i*_(*t*) ∈ [0, 1].

### 2.7 Intraspecies connectedness

Lastly, we define the notion of *intraspecies connectedness*. Essentially, it captures the number of distinct sectors associated with each species that arise via the Voronoi tessellation detailed above. Denote by Γ(*t*) := {*x* ∈ Ω : *t*_FP_(**x**, **m_c_**) = *t*} a circle (in the sense of the front propagation metric *t*_FP_) of radius *t*, centred at **m**_**c**_, which is the centre of mass of the initial patch distribution. Note that the circle lies entirely within the union of the two Voronoi sets 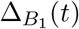 and 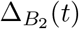, We then classify points on the circle Γ based on which Voronoi set they belong, i.e. we define 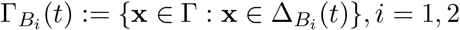. Finally, the *intraspecies connectedness*, *M*, is defined to be half the number points in the intersection between 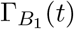 and 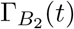, i.e. 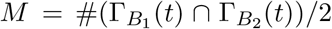. Note that the intersection of 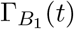 and 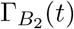 always contains an even, finite number of points due to the definitions of the Voronoi sets 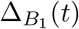 and 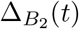 and the circle Γ(*t*).

## 3 Results: Range expansion in spatially heterogeneous domains

With the methods for model analysis in place, we can now determine the effect of spatial landscape heterogeneities. For simplicity, we randomly split the landscape Ω into two types of environment in each independent model realisation (see Section S1 for details). One type represents favourable environmental conditions (large *r*_1_, *r*_2_, *d*_1_, *d*_2_) and the other signifies challenging environmental conditions (small *r*_1_, *r*_2_, *d*_1_, *d*_2_). We assume that the environment is static and remains fixed over the duration of a model simulation. We show in the supplement that results do not depend on this particular choice of heterogeneity by confirming that they hold for other forms of spatially heterogeneous landscapes.

### 3.1 Competition for space

We start by restricting our focus on competition for space only. We do so by considering the dynamics of two differently labelled, but otherwise identical species *B*_1_ and *B*_2_. In terms of the model parameters, this scenario is achieved by setting *d*_1_ = *d*_2_ =: *d, r*_1_ = *r*_2_ =: *r, c*_12_ = *c*_21_ = 0. Unless otherwise stated, we use *k* = 1 throughout and *d* = 0.1, *r* = 5 to characterise favourable environments and *d* = 0.05, *r* = 2.5 to characterise challenging environments. Note the order of magnitude difference between the growth rates and diffusion coefficients. This is to ensure that local growth acts on a faster timescale than spatial spread and results in the width of the expansion front (i.e. the region in which 0 < *B*_1_ + *B*_2_ < *k*) to be small compared to the size of the region behind the front where the population is at carrying capacity. This is a realistic assumption in many contexts, for example the description of cell propagation in bacterial biofilms (Hallatschek et al. 2007), and the migration of sessile organisms (e.g. plants (Deblauwe et al. 2012)) over generational timescales.

#### 3.1.1 Landscape changes cause variability in competitive outcome and area covered by range expansion

We start by considering only changes in the parameter landscape across different model realisations. To cover a wide range of different parameter landscapes, we opt for a Monte Carlo approach in which 500 model realisations with independently chosen parameter landscapes (but fixed initial population distributions with a 1:1 ratio between both species) are performed for each of 16 chosen initial population densities covering three orders of magnitude in initial number of patches (from *N* = 2 to *N* = 500).

As highlighted by the example visualisations shown in Fig. 3.1 for *N* = 6, changes to the spatial landscape cause variability in competitive outcome and area covered by the range expansion. The same trends are observed for other initial population densities within our test range, however, the range of competitive outcome decreases with increasing initial population density and the mean competitive outcome varies across initial population densities (Fig. S5.1).

**Figure 3.1:**
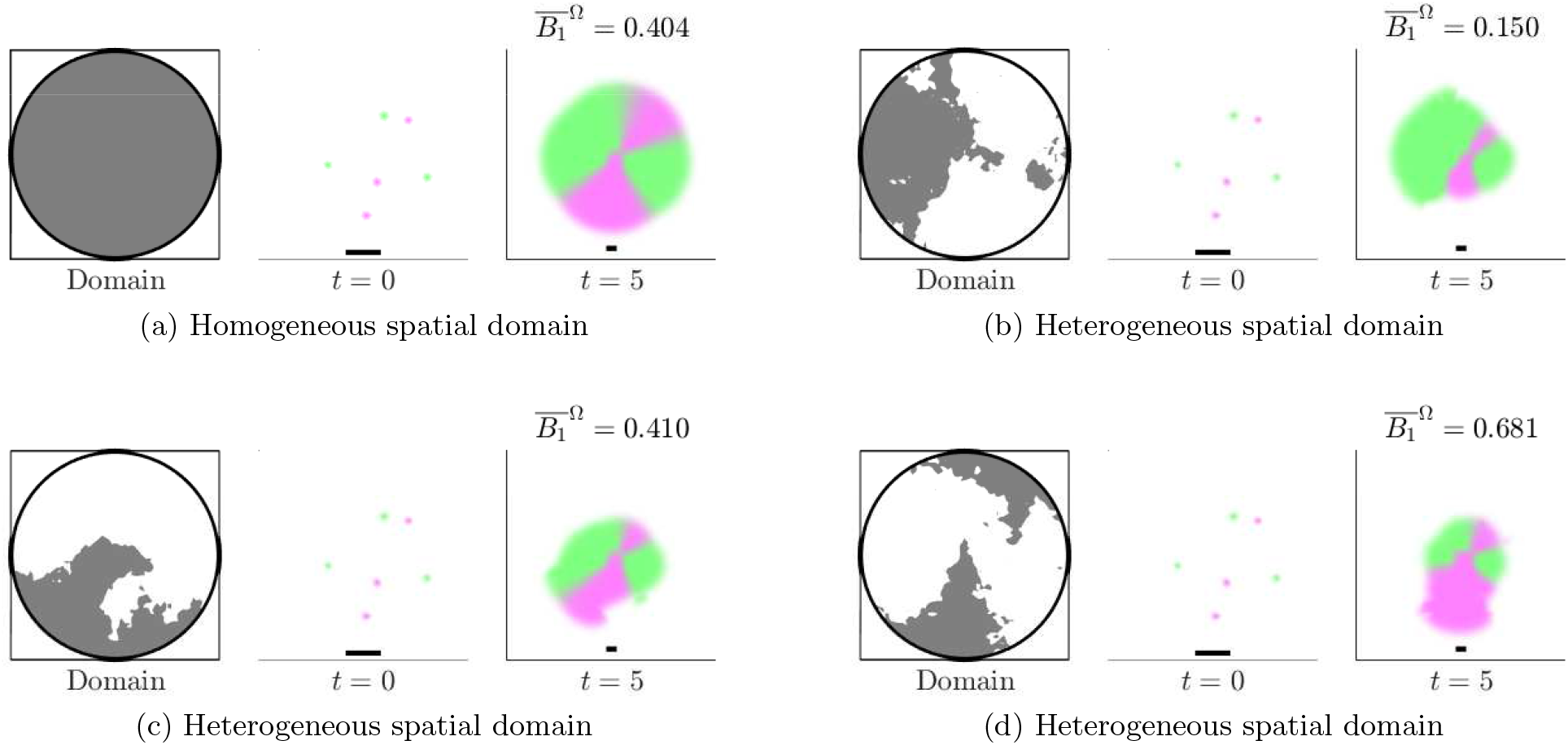
Spatial heterogeneities impact single model realisations. Four model realisations with identical initial conditions but different spatial domains are compared. Part (a) shows a model realisation on a spatially homogeneous domain, part (b-d) on different spatially heterogeneous domains. For each triplet of plots, the left column shows the spatial structure of the domain. Good environmental conditions are shown in grey. The middle column shows the initial condition of the system in a blow-up of the centre of the spatial domain. The right column shows the model result at the defined endpoint. The scale bars are 1 unit length long.

As expected based on experimental literature (Borer et al. 2020; Gralka and Hallatschek 2019), this shows that, all else being fixed, spatial heterogeneities have a significant influence on range expansion in our theoretical framework. Based on this observation, we investigate in the remainder of Section 3.1 (a) whether this source of variability dominates over that induced by randomly chosen initial population distributions; and (b) if predictions of competitive outcome can be made based on the landscape and initial population locations. Finally, in Section 3.2, we investigate (c) what the dominant mode of competition is if species interact antagonistically.

#### 3.1.2 Variability in competitive outcome is a function of initial population density

We next extend our Monte Carlo approach by independently choosing both the parameter landscape and initial population distribution (keeping the initial species ratio fixed at 1:1) at random in each model realisation to assess how simultaneous changes of both properties affect range expansion. In model realisations representing high initial population densities, represented by spatially homogeneous initial conditions, only the parameter landscape is varied.

These simulations reveal that for each fixed initial population density, competitive outcome 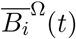 is approximately normally distributed with mean 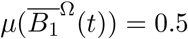. Variability (i.e. standard deviation) in competitive outcome 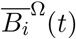 is maximal for 4 ≤ *N* ≤ 20 and decreases with increasing initial population density for *N* > 20 (Fig. 3.2). For high initial population densities, spatial homogeneity is preserved and a 1:1 initial ratio consistently leads to a competitive outcome 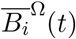 = 0.5, despite the changing parameter landscape across different model realisations (Fig. S5.2). We also note a sharp decrease in variability of competitive outcome if the number of initial patches is decreased to *N* = 2. This also occurs in spatially homogeneous environments and we refer to (Eigentler et al. 2021) for an interpretation of this phenomenon. The reported trends and the observed ranges of competitive outcome are very similar to those reported from spatially homogeneous domains ((Eigentler et al. 2021) and Fig. S5.3). Therefore, we conclude that the initial population density remains the main determinant of variability, despite the landscape changes across different model realisations.

**Figure 3.2:**
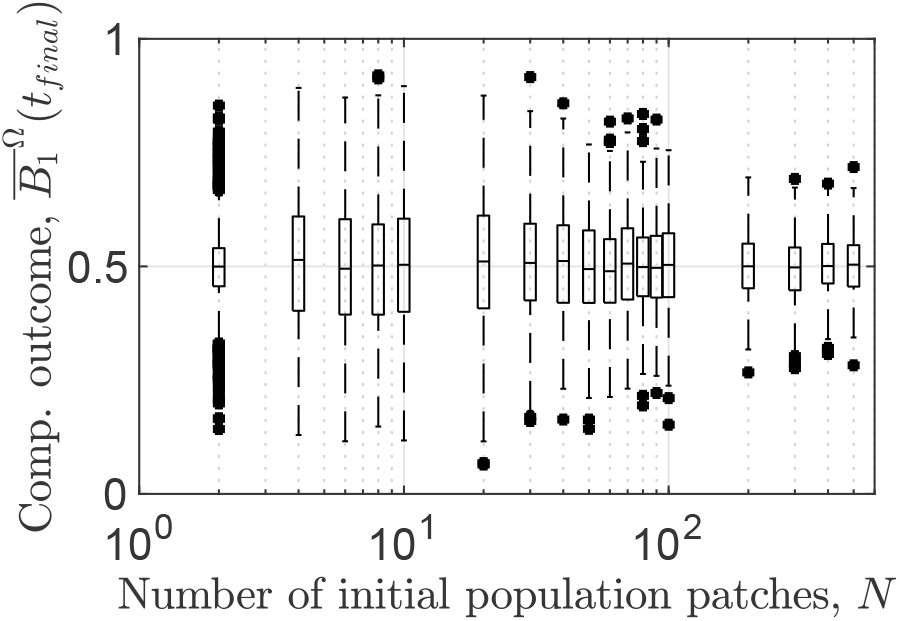
Variability in competitive outcome is a function of initial population density. The relation between initial population density and competitive outcome for the full dataset obtained through our Monte Carlo approach is shown for identical species. For results using different choices of spatially heterogeneous parameter landscapes see Fig. S5.7, Fig. S5.13 and Fig. S5.22.

#### 3.1.3 Variability in area covered by range expansion is determined by spatial heterogeneities

Analysis of data obtained through the Monte Carlo approach further reveals variability in the area covered by range expansion (Fig. 3.3). Moreover, there is a slight increase in area covered by range expansion with increasing initial population density. However, differences in mean area covered between different initial population densities are small compared with the variability for fixed initial population density. Such variability cannot be observed in spatially homogeneous landscapes (Eigentler et al. 2021). Additionally, no variability can be observed either for one of our alternative heterogeneous landscapes (chequerboard pattern, see Section S1) in which the parameter landscape remains unchanged across different model realisations (Fig. S5.8). Moreover, unlike the variability in competitive outcome, the variability in area covered by range expansion is independent of the initial population density (Fig. 3.3). We thus conclude that variability in the area covered by range expansion occurs solely due to changes in the spatially heterogeneous landscapes across different model realisations.

**Figure 3.3:**
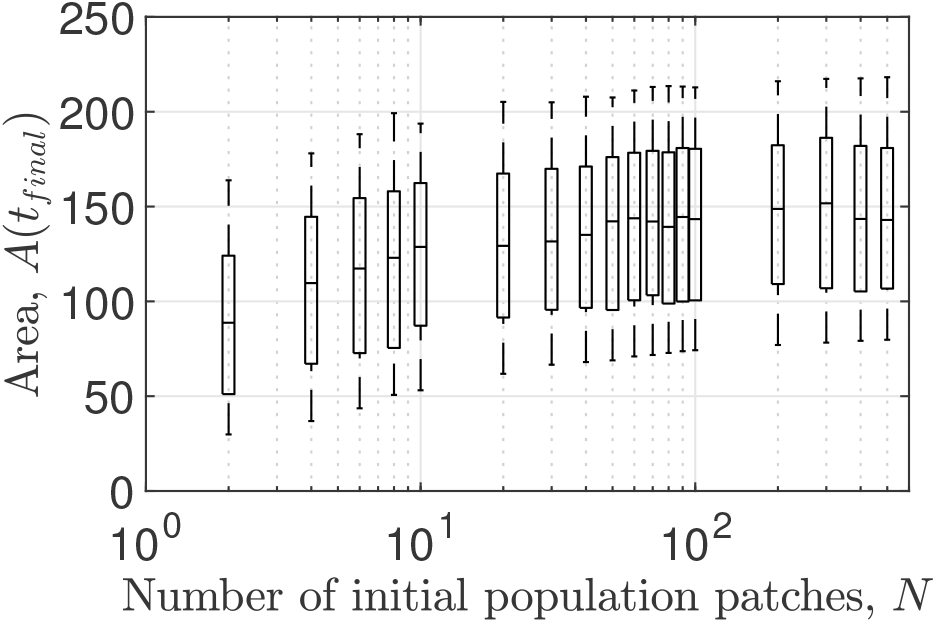
Landscape heterogeneity determines variability in area covered by range expansion. The relation between initial population density and area covered by range expansion is shown for the full dataset obtained through our Monte Carlo approach for identical species. For results using different choices of spatially heterogeneous parameter landscapes see Fig. S5.8, Fig. S5.14 and Fig. S5.20.

#### 3.1.4 Voronoi tessellations predict range expansion, competitive outcome and spatial structure

We now detail how the Voronoi tessellations and the Voronoi index *V*_*i*_(*t*) determine a predictive relationship between the location of the initial patches and the resulting growth dynamics. Using the data from the Monte Carlo approach defined above, the following relationships were established.

First, the Voronoi tessellations provide accurate estimates of the area covered by the population during range expansion (Figs. 3.4b, 3.4d and 3.4f; and Section S3 and Fig. S3.1).

**Figure 3.4:**
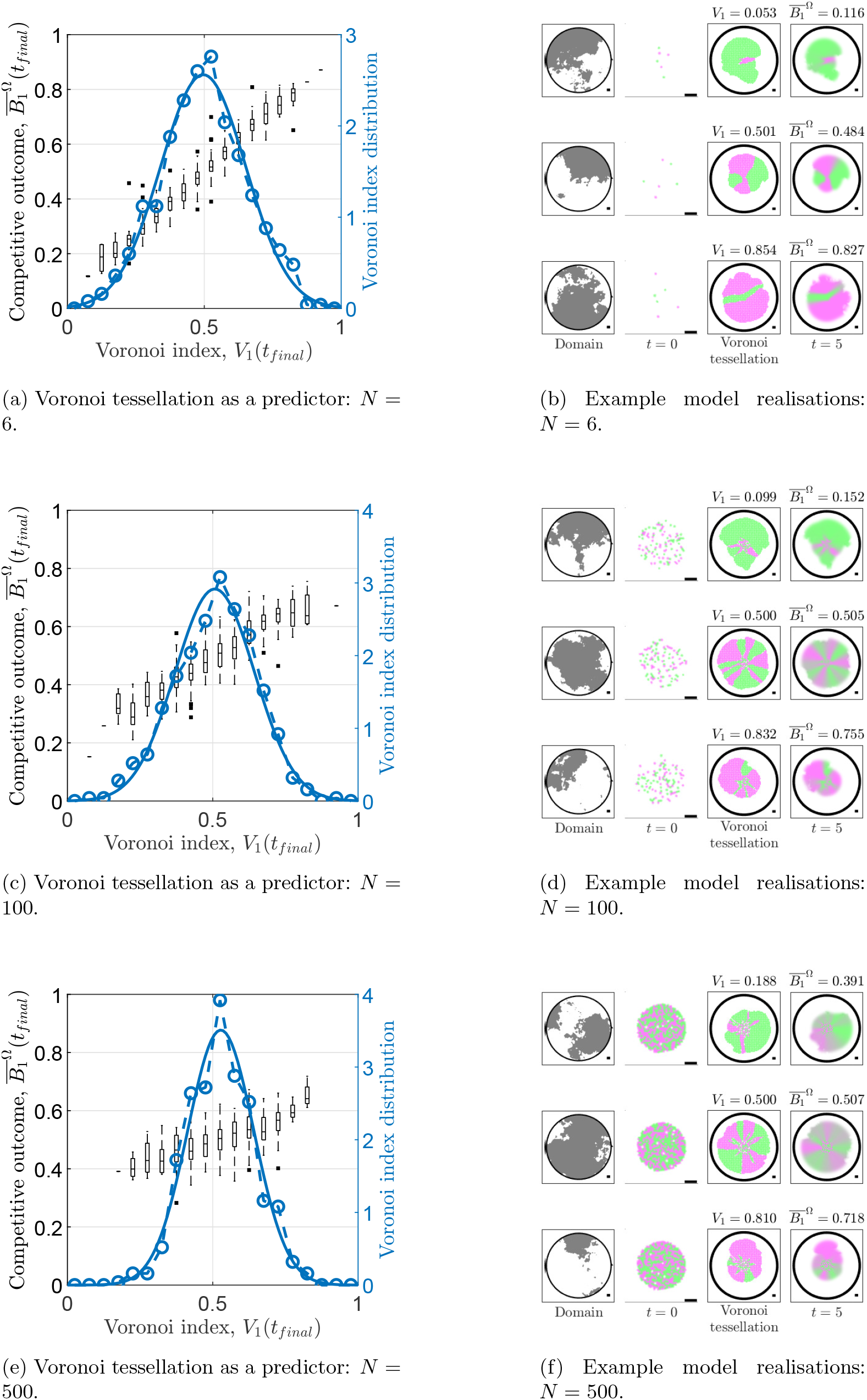
Voronoi tessellations predict competitive outcome. Full figure caption overleaf. Overleaf, the relation between the Voronoi index *V*_1_ and competitive outcome 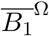 is visualised (black boxplots) for three different initial population densities in (a, c, e). The Voronoi index *V*_1_ is a continuous variable, but for visualisation purposes, the data obtained through our Monte Carlo approach is spit into 20 bins of equal length along the *V*_1_ axis. The distribution of the Voronoi indices is shown as open circles; a fitted normal distribution is shown as a solid line. For each initial population density, example model realisations are shown in (b, d, f). The first column in each of (b, d, f) shows the spatially heterogeneous domain with contour lines; the second column shows the initial condition as a blow-up of Ω_0_; the third column shows the prediction obtained by the Voronoi tessellation and the Voronoi index *V*_1_; and the last column shows the densities at *t* = *t*_final_, the competitive outcome 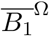 and the spatial assortment *A*. Note that the boundaries between Voronoi sets in spatially homogeneous subsets of the spatial domain do not appear as perfect straight lines because the spatial discretisation of the domain in the calculation of the Voronoi sets. For results using different choices of spatially heterogeneous landscapes see Fig. S5.9, Fig. S5.15 and Fig. S5.21, respectively.

Second, the Voronoi index *V*_*i*_(*t*) acts as an accurate predictor of competitive outcome 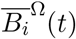 (Fig. 3.4). The accuracy of these predictions is similar to that reported from the spatially homogeneous case for which an alternative prediction method was used (Eigentler et al. 2021). The predictive power of the Voronoi index is time-invariant, even though both the Voronoi index *V*_*i*_(*t*) and the competitive outcome 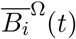 are dynamic quantities that evolve over time during range expansion (Fig. S5.4).

Third, the distributions of the Voronoi indices *V*_*i*_(*t*) are approximately normal with mean *μ*(*V*_*i*_(*t*)) = 0.5 for each fixed initial population density and standard deviation decreasing with initial population density (Figs. 3.4a, 3.4c and 3.4e). Combined with the approximately linear relationship between Voronoi index *V*_*i*_(*t*) and competitive outcome 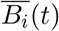, this explains the approximately normal distribution of competitive outcome 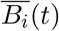.

The results above were obtained for a range of fixed initial population densities, all of which were initiated with a 1 : 1 ratio between species (i.e. 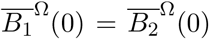 = 0.5). However, the predictive power of the Voronoi index *V*_*i*_(*t*) is robust to variations in initial population ratio as is shown by additional model simulations with fixed initial population density and uniformly randomly chosen initial species ratio (i.e. 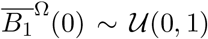) (Fig. S5.5a). While simulations show that increases in initial abundance of a species confers (on average) a competitive advantage, the Voronoi index *V*_*i*_(*t*) provides a more accurate prediction of competitive outcome 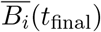) than the initial abundance of the species 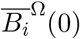 (cf. Fig. S5.5a and Fig. S5.5b).

It is noteworthy that the binary nature of the Voronoi tessellations affects the relation between *V*_*i*_ and competitive outcome 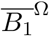. The Voronoi tessellations accurately predict the spatial structure of the model solutions if the initial population density is low (Fig. 3.4b). This further translates to an approximate identity between the Voronoi index *V*_*i*_ and competitive outcome 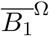 for low initial population densities (i.e. 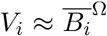). This occurs because species remain spatially segregated during range expansion and only colonise those areas closest to their initial patches. By contrast, the short distances between initial patches at high initial population densities result in more regions of overlap between the the two species in the model solutions. These regions of overlap are not captured by the binary nature of the Voronoi tessellations (Figs. 3.4d and 3.4f) leading to deviations between the Voronoi index *V*_*i*_ and competitive outcome 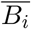. In these cases, changes in the initial patch configuration as measured by the Voronoi index have a smaller impact on competitive outcome (cf. Figs. 3.4a, 3.4c and 3.4e), which leads to the somewhat counter-intuitive conclusion that competition for space between the two species becomes stronger as the initial population becomes more scarce.

### 3.2 Antagonistic interactions

We next consider species that interact antagonistically and investigate the relation between this interaction and spatial dynamics in range expansion. To this end, we set the coefficients in the killing terms to be positive, i.e. *c*_12_ > 0 and *c*_21_ > 0. We assume that species undergo the same growth dynamics in the absence of a competitor species (i.e. *d*_1_ = *d*_2_, *r*_1_ = *r*_2_, but see Sections 4 and S4 for a discussion on relaxation of this assumption) and set *c*_12_ = 5*c*_21_ to create a strong asymmetry between the killing strengths. This assigns *B*_2_ the role of the *intrinsically stronger species* (in the sense of non-spatial dynamics), provided the initial species ratio satisfies *B*_1_(0)*/B*_2_(0) < 5. (Note that there is bistability of both single-species equilibria in the non-spatial system and therefore the definition of the *intrinsically stronger species* depends on both model parameters and initial condition.) Since we typically use *B*_1_(0)*/B*_2_(0) = 1 in our analysis, we refer to *B*_2_ and *B*_1_ as the stronger and weaker species, respectively, throughout this section. Unless otherwise stated, we use the same growth and diffusion parameters as in Section 3.1 and set *c*_21_ = 10, *c*_12_ = 50 and *k* = 1. Note the order of magnitude difference between the killing coefficients and other model parameters. The large size of the killing coefficients ensures that coexistence (in a non-spatial sense) cannot occur as a long transient state.

#### 3.2.1 Antagonistic interactions are the dominant mode of competition for high initial population densities only

We again perform Monte Carlo simulations with fixed initial species ratio but randomly chosen locations of initial population patches and randomly chosen landscapes. Resulting data show that for high initial population density (using both the patch initial conditions with large *N* (Fig. 3.5a) and spatially homogenous initial conditions (not shown)), competitive exclusion of the weaker species occurs consistently. By contrast, coexistence (through spatial segregation) is generally possible for lower initial population densities, but independent model realisations yield variable competitive outcomes (Figs. 3.5b and 3.5c).

**Figure 3.5:**
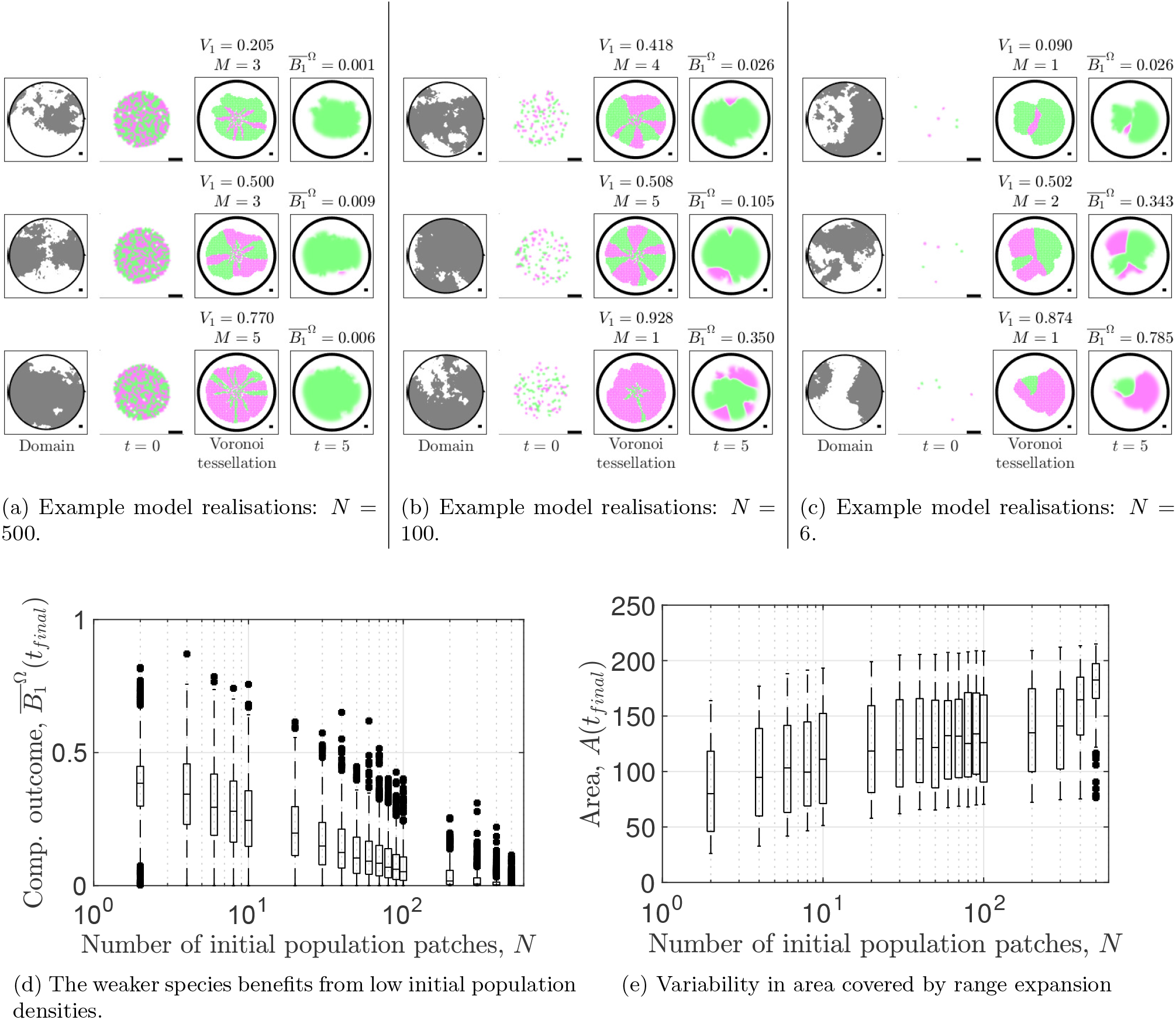
Variability in competitive outcome is a function of initial population density for two antagonistic species. Example model realisations are shown in (a-c) for a selected choice of initial population densities. The first column in each of (a-c) shows the spatially heterogeneous domain with grey areas indicating favourable environments; the second column shows the initial condition as a blow-up of Ω_0_; the third column shows the prediction obtained by the Voronoi tessellation, the Voronoi index *V*_1_ and the intraspecies connectedness *M* ; and the last column shows the densities at *t* = *t*_final_ and 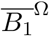. The relation between initial population density and competitive outcome 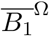 is shown in (d). In (e), the relation between initial population density and area covered by range expansion is shown. For results using different choices of spatially heterogeneous landscapes see Fig. S5.10, Fig. S5.16 and Fig. S5.22 for (d) and Fig. S5.11, Fig. S5.17 and Fig. S5.23 for (e).

Further, similar to the case of identical species, variability in competitive outcome increases with decreasing initial population density (Fig. 3.5d). However, in contrast to the case of identical species, the mean competitive outcome 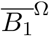 decreases with increasing initial population density *N*. This shows that killing becomes the dominant mode of competition as the initial population density increases. It is important to note that a comparison with data from spatially homogeneous environments highlights that landscape changes across different model realisations do not increase the range of competitive outcomes observed ((Eigentler et al. 2021) and Fig. S5.6).

Finally, similar to the case of identical species, variability in area covered by range expansion is high for all initial population densities (Fig. 3.5e). Mean area covered increases slightly with increasing initial population density, but differences in mean area covered between initial population densities are small compared to the variability observed for fixed initial population density. Again, no such variability occurs in spatially homogeneous landscapes (Eigentler et al. 2021) or if the heterogeneous landscape remains fixed across different model simulations (Fig. S5.11). Thus, we conclude that variability in area covered by range expansion is determined by the landscape changes across different model realisations, even if species interact antagonistically.

#### 3.2.2 Voronoi tessellations predict competitive outcome and highlight that competition for space is the dominant competitive mode for low initial population densities

Similar to the case of identical species, Voronoi tessellations provide an accurate prediction of the area covered by the range expansion (Figs. 3.5a to 3.5c; and Section S3 and Fig. S3.2). Moreover, for each given initial population density, the Voronoi index *V*_1_ is correlated with competitive outcome 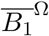 (Fig. 3.6). However, for fixed Voronoi index *V*_1_, high initial population densities yield lower competitive outcomes than low initial population densities on average (cf. Figs. 3.6a and 3.6c). This is in agreement with the dependence of the average competitive outcome on the initial population density shown in Fig. 3.5d. Combined, this highlights that competition for space takes over from antagonisms as the dominant competitive mode as the initial population density decreases.

**Figure 3.6:**
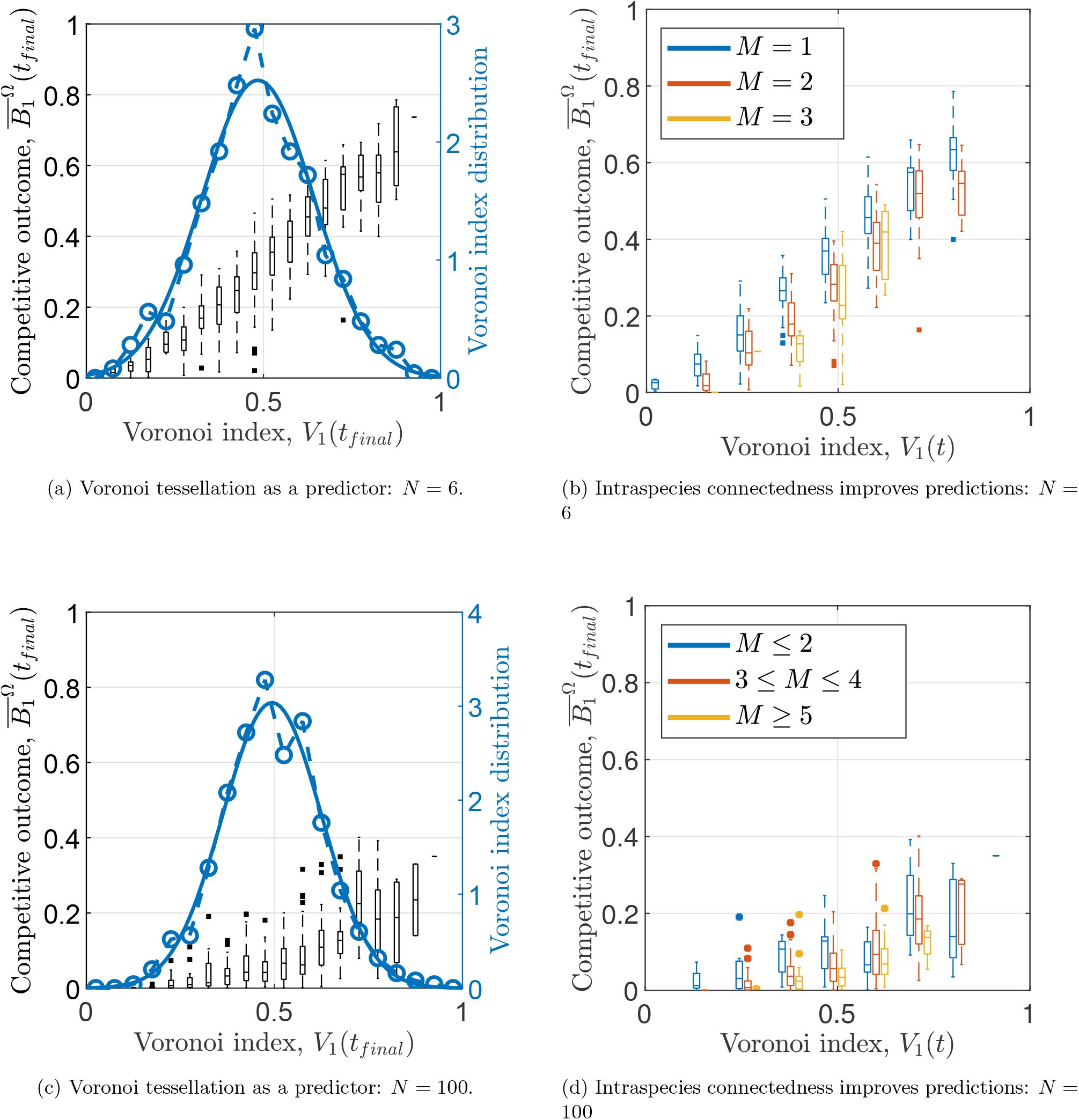
Voronoi tessellations and intraspecies connectedness accurately predict competitive outcome for two antagonistic species. The relation between the Voronoi index *V*_1_ and competitive outcome 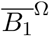 is visualised (black boxplots) for two different initial population densities in (a, c). The Voronoi index *V*_1_ is a continuous variable, but for visualisation purposes, the data obtained through our Monte Carlo approach is spit into 20 bins of equal length along the *V*_1_ axis. The distribution of the Voronoi indices is shown as open circles; a fitted normal distribution is shown as a solid line. In (b, c), the data shown in (a, c) is filtered by the intraspecies connectedness *M*. For visualisation purposes, only 10 bins along the *V*_1_ axis are used. For results using different choices of spatially heterogeneous landscapes see Fig. S5.12, Fig. S5.18 and Fig. S5.24.

#### 3.2.3 Intraspecies connectedness refines predictions of competitive outcome

By definition, the Voronoi index *V*_*i*_ does not account for antagonistic interactions between two species. This leads to a slight decrease in accuracy of the predictions of competitive outcome compared with the case of identical species (cf. Figs. 3.4a and 3.6a). We therefore refine our prediction by taking into account the *intraspecies connectedness*, *M*, (Section 2.7) of the initial population. We hypothesise that more connected initial populations result in less antagonistic interactions during range expansion (essentially, the inter-species boundary is shorter). Indeed, our data show that for fixed Voronoi index *V*_1_, high intraspecies connectedness (low *M*) is beneficial to the weaker species *B*_1_ and leads to a higher competitive outcome 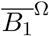 and vice versa. We confirmed that application of the intraspecies connectedness *M* as a filter to the Voronoi index *V*_1_ increases the accuracy of predictions of competitive outcome 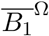, especially if the initial population density is low (Figs. 3.6b and 3.6d).

## 4 Discussion

Previous theoretical and experimental studies have uncovered a wide range of (potentially species-specific) mechanisms underpinning competition during multi-species range expansion (e.g. Aguilera et al. (2019), Burton et al. (2010), Goldschmidt et al. (2017), Legault et al. (2020), Richter et al. (2012) and Weber et al. (2014)). In this paper, we address the impact of spatial heterogeneities in environmental conditions on competition within multi-species range expansion. We reveal that (a) the initial population density is the main determinant of variability in *competitive outcome*; (b) landscape heterogeneities are the main cause of variability in the *area* covered by range expansion; (c) predictions of spatial spread and competitive outcome can be made using a Voronoi tessellation with respect to a suitable metric; (d) for high initial population densities, antagonisms are the dominant mode of competition during range expansion, while for low initial populations densities, the initial distribution determines competitive outcome.

We highlight that during range expansion of identical (but differently labelled) species, the initial species distribution determines competitive outcome (Fig. 3.4). This occurs because the identical species engage in a “race for space” whose spatial structure and global outcome can be predicted accurately by a Voronoi tessellation with respect to an appropriate metric (Figs. 3.4b, 3.4d and 3.4f). This predictor requires as input only information on the spatial structure of the environment, the response of each species to the environment and the initial species distribution. Crucially, no information about the interaction between the species is needed. Indeed, the assumption that descendants colonise areas closest to their ancestors has been experimentally verified (in spatially homogeneous environments only) using microbial species (Lloyd and Allen 2015). While the assumption that competing species are identical is restricting in practice, it presents the ideal test case to reveal the impact of spatial dynamics on competition for related species and forms a basis for further investigation of multi-species range expansion in which other competitive interactions occur.

Our analysis reveals that during multi-species range expansion in which species interactions are subject to antagonistic actions, competition for space remains the dominant mode of competition for low initial population densities, even if the strength of antagonistic actions between species is starkly skewed (Fig. 3.6). This asymmetry prevents the Voronoi index and tessellations predicting spatial structure and competitive outcome with the same degree of accuracy as in the case of identical species (cf. Fig. 3.4b and Fig. 3.5c for example). However, high accuracy can be regained by filtering data with respect to *intraspecies connectedness* - an estimate of the total length of species-to-species interface based on the initial conditions (Figs. 3.6b and 3.6d). The dominance of competition for space over antagonistic mechanisms for low initial population densities is in agreement with modelling studies and field observations that show that traits enhancing dispersal abilities are selected for within population fronts during range expansion (Phillips et al. 2008; Travis and Dytham 2002).

Model analysis highlights that species coexistence can occur through spatial segregation with limited overlap between the species along species-to-species interfaces if species interact through antagonistic actions. Therefore, spatial segregation offers protection from competitors. In particular, the majority of the population within the expansion front is unaffected by the antagonistic actions. This enables an intrinsically weaker species to coexist with a stronger competitor, provided the former is able to spatially segregate from its competitor in the early stages of range expansion. This highlights that classical mechanisms commonly commonly associated with enabling coexistence, such as a trade-off between local competitiveness and dispersal abilities (e.g. Gravel et al. (2010), Hassell et al. (1994), Horn and MacArthur (1972), Levins and Culver (1971) and Tilman (1994)) or non-transitive competitive hierarchies (“rock-paper-scissors”) (e.g. Avelino et al. (2019), Kerr et al. (2002), Lowery and Ursell (2019) and Reichenbach et al. (2007)) are not necessarily required for coexistence to occur. Instead, our results show that spatial dynamics alone are sufficient to generate coexistence in range expansion originating from low initial population densities.

The model revealed that, *on average*, spatially heterogeneous landscapes do not affect competitive outcome. However, outcomes from single realisations can be affected greatly by domain heterogeneities (Fig. 3.1). Moreover, variability in the area covered by range expansion is induced by spatial heterogeneities, independent of the initial population density (Figs. 3.3 and 3.5e). These results highlight the importance of considering landscape heterogeneities when implementing human-driven interventions in ecological systems. Our results predict that in cases where large numbers of organisms are introduced and undergo range expansion simultaneously (e.g. in the addition of biofertilizers to soil (Arroyave-Toro et al. 2017; Calvo-Garrido et al. 2019)), spatial landscape heterogeneities would be rendered insignificant. By contrast, landscape heterogeneities and initial population distributions would need to be carefully considered in applications for which only a small number of individuals are added into a system (e.g. rewilding of mammals (Lorimer et al. 2015)).

We focussed our analysis on a generic competition model in which dynamics of species in the absence of interspecific competition are governed by logistic growth and diffusion. However, we argue that our method could be extended to species whose growth and dispersal behaviour is governed by other functional responses. Generalisation would only require the calculation/estimation of the front propagation speed *c*(**x**) used in the definition of the front propagation metric *t*_FP_. If front expansion dynamics cannot be approximated by *pulled* travelling waves, the front expansion speed *c*(**x**) may additionally depend on other model parameters such as the population’s carrying capacity or intraspecific competition coefficients (which may also be space-dependent). Therefore, calculation of an estimate of *c*(**x**) may not be as straightforward as for the model considered in this paper, but could nevertheless be approximated by numerical or field experiments. Moreover, our method could be generalised to two or more species whose growth and dispersal dynamics differ from each other. In such a case, the front expansion speed *c*(**x**) would differ for each species and therefore would require the definition of separate metrics. With a slight redefinition of the Voronoi sets, the definition of the Voronoi index remains unchanged. For a mathematically rigorous definition of the Voronoi indices in this case, see Section S4.

## 5 Conclusion

From the human perspective, range expansion is becoming an increasingly important process across many different spatial and temporal scales. Its significance extends from applications of fungal species and genetically modified, biofilm-forming microbes in biological, medical and industrial settings (Dzianach et al. 2019; Martignoni et al. 2020) to conservation programs for plant and animal species whose habitats are shifting polewards (or towards higher altitudes) due to climate change (Rosenzweig et al. 2007). Our results reveal that traits required for competitive success during range expansion starkly differ from those characterising competitive fitness in equilibrium settings if range expansion originates from low population density. This shows that intrinsically weaker species are able to persist (or in rare cases even outcompete) stronger species during range expansion, provided they are able to perform well in the “race for space” that determines competitive outcome. Thereby, our results provide a complement to field studies and evolutionary models that have shown dispersal traits within single species are selected for within the expansion front (Phillips et al. 2008; Travis and Dytham 2002). While our theoretical framework is deliberately kept simple so as to be applicable in a general setting, the model and methods form a foundation for extensions and applications to specific ecosystems. Adaptations of the framework to specific species in specific environments would require field data on an ecosystem-wide scale. Such data is becoming increasingly available thanks to advances in remote sensing technologies (e.g. Deblauwe et al. (2012)) and machine learning applications (e.g. Lary et al. (2016)) which highlights the potential of the theoretical framework to support future work.

## Data availability statement

Computational code associated with this study is available on Github and has been archived by Zenodo (Eigentler 2021).

## Acknowledgements

Work presented in this paper was funded by the Biotechnology and Biological Science Research Council (BBSRC) [BB/P001335/1, BB/R012415/1]. We are grateful to members of the Stanley-Wall lab and Prof Cait E. MacPhee (University of Edinburgh) for helpful discussions.

## Author contributions

Lukas Eigentler: Conceptualization, Methodology, Software, Validation, Formal analysis, Data Curation, Writing - Original Draft, Visualization, Project administration Nicola R. Stanley-Wall: Conceptualization, Writing - Review & Editing, Supervision, Project administration, Funding acquisition

Fordyce A. Davidson: Conceptualization, Methodology, Writing - Original Draft, Writing - Review & Editing, Supervision, Project administration, Funding acquisition

