## Supplementary material for "A theoretical framework for multi-species range expansion in spatially heterogeneous landscapes"

Lukas Eigentler[1,2,*], Nicola R. Stanley-Wall[1], Fordyce A. Davidson[2,*]

[1]Division of Molecular Microbiology, School of Life Sciences, University of Dundee, Dundee DD1 5EH, United Kingdom
[2]Division of Mathematics, School of Science and Engineering, University of Dundee, Dundee DD1 4HN, United

### Contents

**S1 Spatial heterogeneity** **S2**
S1.1 Generation of a fractal-like random surface . . . . . . . . . . . . . . . . . . . . . . . . S3

**S2 Proof that $t_{\mathrm{FP}}(\mathbf{x}, \mathbf{y})$ is a metric** **S3**

**S3 Accuracy of predictions of area covered by range expansion** **S6**

**S4 Voronoi index for systems of two or more species with different front expansion speeds** **S6**

**S5 Supplementary figures** **S10**

### List of Figures

S1.1 Spatially heterogeneous parameter landscapes. . . . . . . . . . . . . . . . . . . . . . . . S4
S1.2 Spatially heterogeneous landscape. . . . . . . . . . . . . . . . . . . . . . . . . . . . . . . S5
S3.1 Accuracy of predictions of area covered for identical species. . . . . . . . . . . . . . . . . S7
S3.2 Accuracy of predictions of area covered for antagonistic species. . . . . . . . . . . . . . S8
S5.1 Changes in spatial landscape induce variability in competitive outcome and area covered by range expansion if the initial population distribution is fixed. . . . . . . . . . . . . S10
S5.2 Spatial homogeneity is preserved in range expansion of identical species. . . . . . . . . S11
S5.3 Initial population density is the main determinant of variability in competitive outcome for identical species. . . . . . . . . . . . . . . . . . . . . . . . . . . . . . . . . . . . . S12
S5.4 Voronoi indices predict competitive outcome at all times. . . . . . . . . . . . . . . . . S12
S5.5 Voronoi index determines competitive outcome independent of initial population ratio. S13
S5.6 Initial population density is the main determinant of variability in competitive outcome for antagonistic species. . . . . . . . . . . . . . . . . . . . . . . . . . . . . . . . . . . . S13
S5.7 Competitive outcome for two identical species as a function of initial population density in heterogeneous landscapes case (i). . . . . . . . . . . . . . . . . . . . . . . . . . . . S14
S5.8 Spatial heterogeneities determine variability in area covered by range expansion in heterogeneous landscapes case (i). . . . . . . . . . . . . . . . . . . . . . . . . . . . . . . . S14
S5.9 Voronoi index as a predictor of competitive outcome for two identical species in heterogeneous landscapes case (i). . . . . . . . . . . . . . . . . . . . . . . . . . . . . . . . . . S15
S5.10 Competitive outcome of two antagonistic species as a function of initial population density in heterogeneous landscapes case (i). . . . . . . . . . . . . . . . . . . . . . . . . . S16

S5.11 Spatial heterogeneities determine variability in area covered by range expansion of antagonistic species in heterogeneous landscapes case (i). . . . . . S16
S5.12 Voronoi index as a predictor of competitive outcome for two antagonistic species in heterogeneous landscapes case (i). . . . . . S17
S5.13 Competitive outcome for two identical species as a function of initial population density in heterogeneous landscapes case (ii). . . . . . S18
S5.14 Spatial heterogeneities determine variability in area covered by range expansion in heterogeneous landscapes case (ii). . . . . . S18
S5.15 Voronoi index as a predictor of competitive outcome for two identical species in heterogeneous landscapes case (ii). . . . . . S19
S5.16 Competitive outcome of two antagonistic species as a function of initial population density in heterogeneous landscapes case (ii). . . . . . S20
S5.17 Spatial heterogeneities determine variability in area covered by range expansion of antagonistic species in heterogeneous landscapes case (ii). . . . . . S20
S5.18 Voronoi index as a predictor of competitive outcome for two antagonistic species in heterogeneous landscapes case (ii). . . . . . S21
S5.19 Competitive outcome for two identical species as a function of initial population density in heterogeneous landscapes case (iv). . . . . . S22
S5.20 Spatial heterogeneities determine variability in area covered by range expansion in heterogeneous landscapes case (iv). . . . . . S22
S5.21 Voronoi index as a predictor of competitive outcome for two identical species in heterogeneous landscapes case (iv). . . . . . S23
S5.22 Competitive outcome of two antagonistic species as a function of initial population density in heterogeneous landscapes case (iv). . . . . . S24
S5.23 Spatial heterogeneities determine variability in area covered by range expansion of antagonistic species in heterogeneous landscapes case (iv). . . . . . S24
S5.24 Voronoi index as a predictor of competitive outcome for two antagonistic species in heterogeneous landscapes case (iv). . . . . . S25

### S1 Spatial heterogeneity

We account for spatial heterogeneity in the computational domain $\Omega$ by employing space-dependent diffusion coefficients and growth rates, i.e. $d_1 = d_1(\mathbf{x})$, $d_2 = d_2(\mathbf{x})$, $r_1 = r_1(\mathbf{x})$ and $r_2 = r_2(\mathbf{x})$. These heterogeneities represent variations in environmental conditions, such as nutrient concentration or availability of nesting sites in a generic way. We generate a "landscape" of diffusion coefficients and growth rates by linking them to a surface $L(\mathbf{x})$. There exist countless options to define $L$. We use a range of different test cases that range from periodic domains (case (i)) to randomly chosen parameter landscapes with a fractal-like structure (case (iii) and (iv)):

Case (i): We define a periodic parameter landscape by letting $L$ alternate between $L = 1$ and $L = -1$ in a chequerboard pattern and setting $r_i = r_i^{\max}$ if $L = 1$ and $r_i = r_i^{\max}/2$ otherwise and $d_i = d_i^{\max}$ if $L = 1$ and $d_i = d_i^{\max}/2$ otherwise (Fig. S1.1a).

Case (ii): We introduce randomness to the landscape by utilising the same spatial grid as in case (i), but choosing between $L = 1$ and $L = -1$ uniformly at random and setting model parameters as above (Fig. S1.1b).

Case (iii): Cases (i) and (ii) create a parameter landscape in which randomness occurs at a fixed spatial scale, defined by the size of elements in the spatial grid. To take into account randomness across different scales, we define $L$ to be a fractal-like surface, generated by a recursive algorithm (see Chen and Yang 2016; Fournier et al. 1982 and Section S1.1). Note that $L$ is not a true fractal because the recursive algorithm is stopped after a small finite number of iterations. Termination guarantees that the randomness in $L$ occurs at a spatial scale no finer than that of the spatial mesh to solve the model equations. In this case, we utilise $L$ to generate a parameter landscape by setting $r_i = r_i^{\max}$ if $L \geq 0$ and $r_i = r_i^{\max}/2$ otherwise and $d_i = d_i^{\max}$ if $L \geq 0$ and $d_i = d_i^{\max}/2$ otherwise (Fig. S1.1c).

Case (iv): In the final test case, the fractal-like surface $L$ (see case (iii)) is used to define a continuous landscape of diffusion coefficients and growth rates in which large values of $L$ are associated with better environmental conditions than small values of $L$. This is done by setting $r_i = r^{\text{scale}}(r_i^{\text{mean}} + L)$ if $L > -r_i^{\text{mean}}$ and $r_i = 0$ if $L \leq -r_i^{\text{mean}}$ and $d_i = d^{\text{scale}}(d_i^{\text{mean}} + L)$ if $L > -d_i^{\text{mean}}$ and $d_i = 0$ if $L \leq -d_i^{\text{mean}}$ (Fig. S1.1d).

In the main text, we only present results for case (iii). However, we show in the supplementary material that results are not case-specific.

#### S1.1 Generation of a fractal-like random surface

In cases (iii) and (iv), a "landscape" of diffusion coefficients and growth rates is generated by linking them to a random surface $L$ with monofractal structure, generated using the random midpoint displacement method Chen and Yang 2016; Fournier et al. 1982. Briefly, the recursive algorithm constructs a surface as follows (see Chen and Yang 2016 for full details). As a preliminary step, values drawn from the standard normal distribution $\mathcal{N}(0, 1)$ are assigned to the corners of the smallest square enclosing the computational domain $\Omega$. Linear interpolation creates a two-dimensional surface $L_0$ in $\mathbb{R}^3$ consisting of one quadrilateral element (Fig. S1.2(a)). Then, in the first $(n = 1)$ step of the recursion, random perturbations drawn from $\mathcal{N}(0, \sigma_n^2)$, where $\sigma_n^2 = 2^{-2H(n+1)}$, $0 \leq H \leq 1$, are added to the centre of the quadrilateral and all bisection points of edges connecting the corners of the quadrilateral. Linear interpolation then creates a surface $L_1$ consisting of four quadrilaterals (Fig. S1.2(b)). In consequent steps of the recursion, these steps are repeated for all quadrilaterals comprising the surface until the required spatial resolution is reached after $N$ steps (Fig. S1.2(c-g)). Here, we stop after the maximum number of iterations that result in a surface described by less points than the number of nodes in the spatial mesh used in the application of a finite element method to solve the model. This ensures that the spatial landscape is described on a coarser scale than the spatial mesh and thus enables the finite element method to capture the whole complexity of the spatial heterogeneity. The coefficient $H$ is referred to as the Hurst exponent and determines the fractal dimension $d_{\text{frac}}$ of the surface as $d_{\text{frac}} = 3 - H$ Chen and Yang 2016. The fractal dimension $d_{\text{frac}}$ of a surface characterises how the number of cubes with sidelength $a$ required to fully cover the surface, denoted by $N(a)$, changes with the sidelength $a$ as $N(a) \propto (1/a)^{d_{\text{frac}}}$. Thus, a surface with fractal dimension $d_{\text{frac}} \approx 2$ fills space like a non-fractal surface, while a surface with $d_{\text{frac}} \approx 3$ fills space like a volume. Here, we use $H = 0.5$ resulting in a fractal dimension of $d_{\text{frac}} = 2.5$.

### S2 Proof that $t_{\text{FP}}(\mathbf{x}, \mathbf{y})$ is a metric

**Proposition S2.1** *The function $t_{\text{FP}}(\mathbf{x}, \mathbf{y})$ is a metric.*

#### Proof

(i) (identity of indiscernibles) If $\mathbf{x} = \mathbf{y}$, then there exists a path $P \in \mathcal{P}(\mathbf{x}, \mathbf{y})$ parameterised by $p(\tau) \equiv \mathbf{x}$. Then $I(P) = \int_0^1 \frac{1}{c(p(\tau))} \|p'(\tau)\| \mathrm{d}\tau = 0$, because $\|p'(\tau)\| \equiv 0$. Hence, also $t_{\text{FP}}(\mathbf{x}, \mathbf{y}) = 0$.

(ii) (identity of indiscernibles cont.) If $t_{\text{FP}}(\mathbf{x}, \mathbf{y}) = 0$, then there exists a path $P \in \mathcal{P}(\mathbf{x}, \mathbf{y})$ such that $I(P) = \int_0^1 \frac{1}{c(p(\tau))} \|p'(\tau)\| \mathrm{d}\tau = 0$. Since $c(p(\tau)) < \infty$ for all $0 \leq \tau \leq 1$, this yields $\|p'(\tau)\| \equiv 0$. The operator $\|\cdot\|$ is a norm and thus $p'(\tau) \equiv 0$ for all $\tau$. Therefore $p(\tau) \equiv$ const., and in particular $p(0) = p(1)$, which gives $\mathbf{x} = \mathbf{y}$.

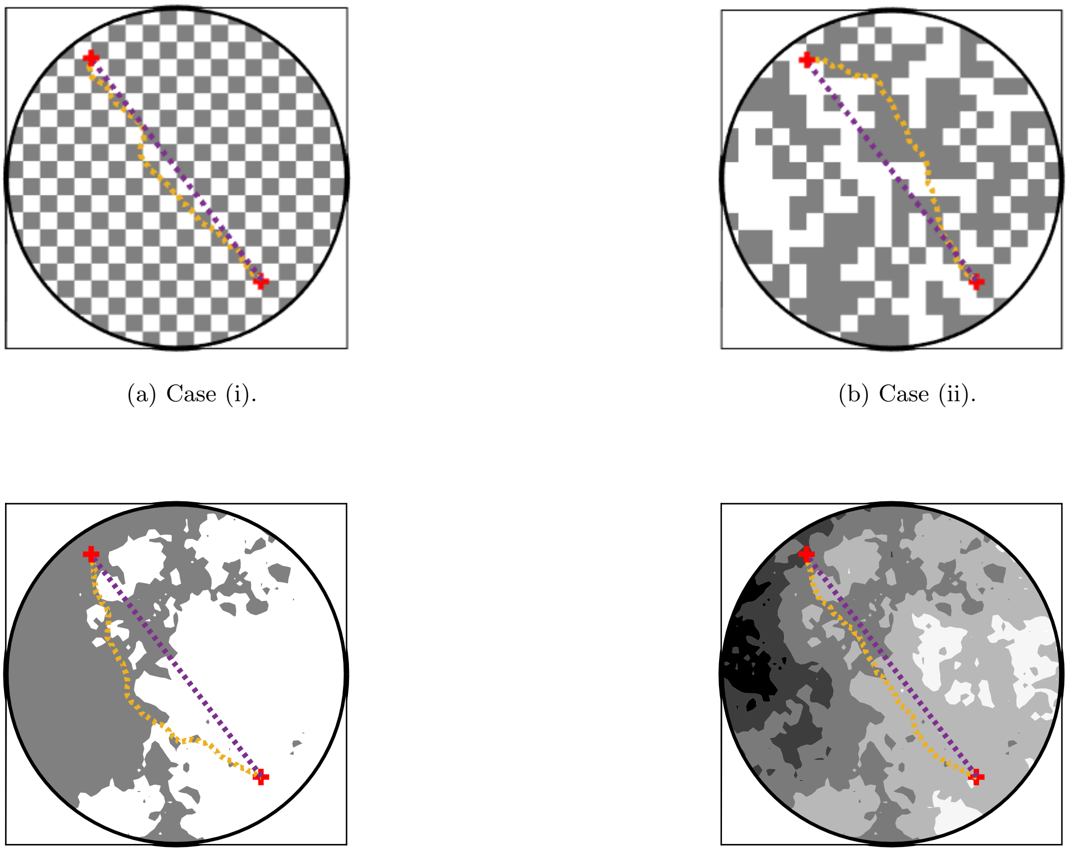

(a) Case (i). (b) Case (ii).

(c) Case (iii). (d) Case (iv).

Figure S1.1: **Spatially heterogeneous parameter landscapes.** In (a), a periodic chequerboard landscape distinguishing between two different parameter regimes is shown (case (i)). In (b), the chequerboard pattern is changed by choosing between the two parameter regimes uniformly at random in each box (case (ii)). In (c), the monofractal is split into two regions by the contour $L = 0$ to create a heterogeneous parameter landscape according to case (iii). In (d), the monofractal is used to defined a continuous parameter landscape according to case (iv). In all plots, the shortest path in the sense of the front propagation metric $t_{\mathrm{FP}}$ (yellow) is compared with the shortest Euclidean distance (purple) between two points.

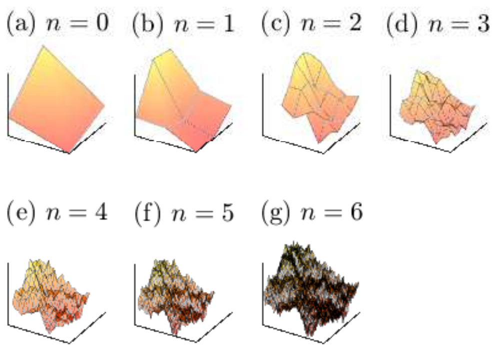


Recursive generation of a monofractal surface.

Figure S1.2: **Spatially heterogeneous landscape.** The recursive generation of a monofractal surface is shown in (a-g). For details on the recursion see Section S1.1.

(iii) (symmetry) If $P \in \mathcal{P}(\mathbf{x},\mathbf{y})$ is parameterised by $p(\tau)$, then the path $Q$, parameterised by $q(\tau) = p(1-\tau)$ is a member of $\mathcal{P}(\mathbf{y},\mathbf{x})$. Then,

$$t_{\mathrm{FP}}(\mathbf{y},\mathbf{x}) = \inf_{Q\in\mathcal{P}(\mathbf{y},\mathbf{x})} I(Q) = \inf_{Q\in\mathcal{P}(\mathbf{y},\mathbf{x})} \left[\int_0^1 \frac{1}{c(q(\tau))}\|q'(\tau)\|\mathrm{d}\tau\right]$$
$$= \inf_{P\in\mathcal{P}(\mathbf{x},\mathbf{y})} \left[\int_0^1 \frac{1}{c(p(1-\tau))}\|p'(1-\tau)\|\mathrm{d}\tau\right].$$

The change of coordinates $\tilde{\tau} = 1-\tau$ gives

$$t_{\mathrm{FP}}(\mathbf{y},\mathbf{x}) = \inf_{P\in\mathcal{P}(\mathbf{x},\mathbf{y})} \left[-\int_1^0 \frac{1}{c(p(\tilde{\tau}))}\|p'(\tilde{\tau})\|\mathrm{d}\tilde{\tau}\right] = \inf_{P\in\mathcal{P}(\mathbf{x},\mathbf{y})} \left[\int_0^1 \frac{1}{c(p(\tilde{\tau}))}\|p'(\tilde{\tau})\|\mathrm{d}\tilde{\tau}\right] =$$
$$\inf_{P\in\mathcal{P}(\mathbf{x},\mathbf{y})} I(P) = t_{\mathrm{FP}}(\mathbf{x},\mathbf{y}).$$

(iv) (triangle inequality) Let $P_1 \in \mathcal{P}(\mathbf{x},\mathbf{y})$ be parameterised by $p_1$ and $P_2 \in \mathcal{P}(\mathbf{y},\mathbf{z})$ be parameterised by $p_2$ such that $t_{\mathrm{FP}}(\mathbf{x},\mathbf{y}) = \inf_{P\in\mathcal{P}(\mathbf{x},\mathbf{y})} I(P) = I(P_1)$ and $t_{\mathrm{FP}}(\mathbf{y},\mathbf{z}) = \inf_{P\in\mathcal{P}(\mathbf{y},\mathbf{z})} I(P) = I(P_2)$. Using the parameterisations, we define new functions $q_1 : [0,1/2] \to \Omega : \tau \mapsto p_1(2\tau)$, $q_2 : [1/2,1] \to \Omega : \tau \mapsto p_2(2\tau-1)$ and

$$q : [0,1] \to \Omega : \tau \mapsto \begin{cases} q_1(\tau) & \text{if} \quad 0 \le \tau \le \frac{1}{2} \\ q_2(\tau) & \text{if} \quad \frac{1}{2} \le \tau \le 1 \end{cases}.$$

The function $q$ is a parameterisation of a path $Q \in \mathcal{P}(\mathbf{x},\mathbf{z})$, because $q(0) = q_1(0) = p_1(0) = \mathbf{x}$, $q(1) = q_2(1) = p_2(1) = \mathbf{z}$, $q_1(1/2) = p_1(1) = \mathbf{y} = p_2(0) = q_2(1/2)$ and $q$ is continuously differentiable almost everywhere. Then

$$I(Q) = \int_0^1 \frac{1}{c(q(\tau))}\|q'(\tau)\|\mathrm{d}\tau = \int_0^{\frac{1}{2}} \frac{1}{c(q_1(\tau))}\|q_1'(\tau)\|\mathrm{d}\tau + \int_{\frac{1}{2}}^1 \frac{1}{c(q_2(\tau))}\|q_2'(\tau)\|\mathrm{d}\tau =$$
$$\int_0^{\frac{1}{2}} \frac{1}{c(p_1(2\tau))}\|p_1'(2\tau)\|\mathrm{d}\tau + \int_{\frac{1}{2}}^1 \frac{1}{c(p_2(2\tau-1))}\|p_2'(2\tau-1)\|\mathrm{d}\tau.$$

The changes of variables $\kappa = 2\tau$ and $\kappa = 2\tau - 1$ in the first and second integral, respectively, give

$$I(Q) = \int_0^1 \frac{1}{c(p_1(\kappa))} \|p_1'(\kappa)\| \mathrm{d}\kappa + \int_0^1 \frac{1}{p_2(\kappa)} \|p_2'(\kappa)\| \mathrm{d}\kappa$$
$$= I(P_1) + I(P_2) = t_{\mathrm{FP}}(\mathbf{x}, \mathbf{y}) + t_{\mathrm{FP}}(\mathbf{y}, \mathbf{z}).$$

Therefore

$$t_{\mathrm{FP}}(\mathbf{x}, \mathbf{z}) = \inf_{P \in \mathcal{P}(\mathbf{x}, \mathbf{z})} I(P) \leq I(Q) = t_{\mathrm{FP}}(\mathbf{x}, \mathbf{y}) + t_{\mathrm{FP}}(\mathbf{y}, \mathbf{z}).$$

### S3 Accuracy of predictions of area covered by range expansion

We quantify the accuracy of estimates of the area covered by range expansion as follows. The area estimate given by the Voronoi tessellation is the area covered by $\Delta_{B_1}(t) \cup \Delta_{B_2}(t)$. We define the area covered by range expansion in our model simulation as the area of the set $\{x \in \Omega : B_1(t, x) + B_2(t, x) > 0.1k_{\mathrm{cap}}\}$. To compare the estimate with the model realisation, we consider a grid of equidistant points covering the computational domain $\Omega$. We define two vectors, $A_{\mathrm{pred}}$ and $A_{\mathrm{sim}}$, as follows. For the i-th element $x_i$ of the grid covering the domain, we set the i-th element of $A_{\mathrm{pred}} = 1$ if $x_i \in \Delta_{B_1}(t) \cup \Delta_{B_2}(t)$, and $A_{\mathrm{pred}} = 0$ otherwise. Similarly, we set $A_{\mathrm{sim}} = 1$ if $x_i \in \{x \in \Omega : B_1(t, x) + B_2(t, x) > 0.1k_{\mathrm{cap}}\}$ and $A_{\mathrm{sim}} = 0$ otherwise. This creates two binary vectors $A_{\mathrm{pred}}$ and $A_{\mathrm{sim}}$ quantifying which points in the computational domain lie within the predicted region of range expansion and the realised region of range expansion in model simulations, respectively. We then quantify the similarity between both vectors by calculating the linear correlation between $A_{\mathrm{pred}}$ and $A_{\mathrm{sim}}$.

Calculation of linear correlation between $A_{\mathrm{pred}}$ and $A_{\mathrm{sim}}$ for our data set obtained from Monte Carlo simulations (see main text) reveals that correlation is high for all landscape cases (with a slight reduction in accuracy for case (i)) and all initial population densities for both identical (Fig. S3.1) and antagonistic (Fig. S3.2) species.

### S4 Voronoi index for systems of two or more species with different front expansion speeds

Assume there are $K \in \mathbb{N}$ different species $B_i, i = 1, \ldots, K$, whose front propagation speeds (in the absence of competition) $c_i(\mathbf{x})$ differ from each other. Then, the front propagation metric for species $B_i$, denoted by $t_{\mathrm{FP}}^{(i)}$, is defined by

$$t_{\mathrm{FP}}^{(i)}(\mathbf{x}, \mathbf{y}) := \inf_{P \in \mathcal{P}(\mathbf{x}, \mathbf{y})} \int_P \frac{1}{c_i(\mathbf{x})}, \quad i = 1, \ldots, K.$$

Denote the $N_i \in \mathbb{N}$ initial patch locations of species $B_i$ by $\mathcal{B}_i := \{\mathbf{x}_1^{(i)}, \ldots, \mathbf{x}_{N_i}^{(i)}\}, i = 1, \ldots K$. Then, the Voronoi set for the initial patch $\mathbf{x}_j^{(i)}, j = 1, \ldots, N_i$, is given by

$$\Delta_{i,j}^{\Omega} := \left\{ \mathbf{x} \in \Omega : t_{\mathrm{FP}}^{(i)}(\mathbf{x}, \mathbf{x_j^{(i)}}) \leq \min_{m=1,\ldots,N_\ell} t_{\mathrm{FP}}^{(\ell)}(\mathbf{x}, \mathbf{x_m^{(\ell)}}) \quad \text{for all } \ell = 1, \ldots, K, \ell \neq i \right\},$$
$$i = 1, \ldots, K, j = 1, \ldots, N_i.$$

Then, the Voronoi set for species $B_i$ is

$$\Delta_{B_i}^{\Omega} := \bigcup_{j=1,\ldots,N_i} \Delta_{i,j}^{\Omega}, \quad i = 1, \ldots, K.$$

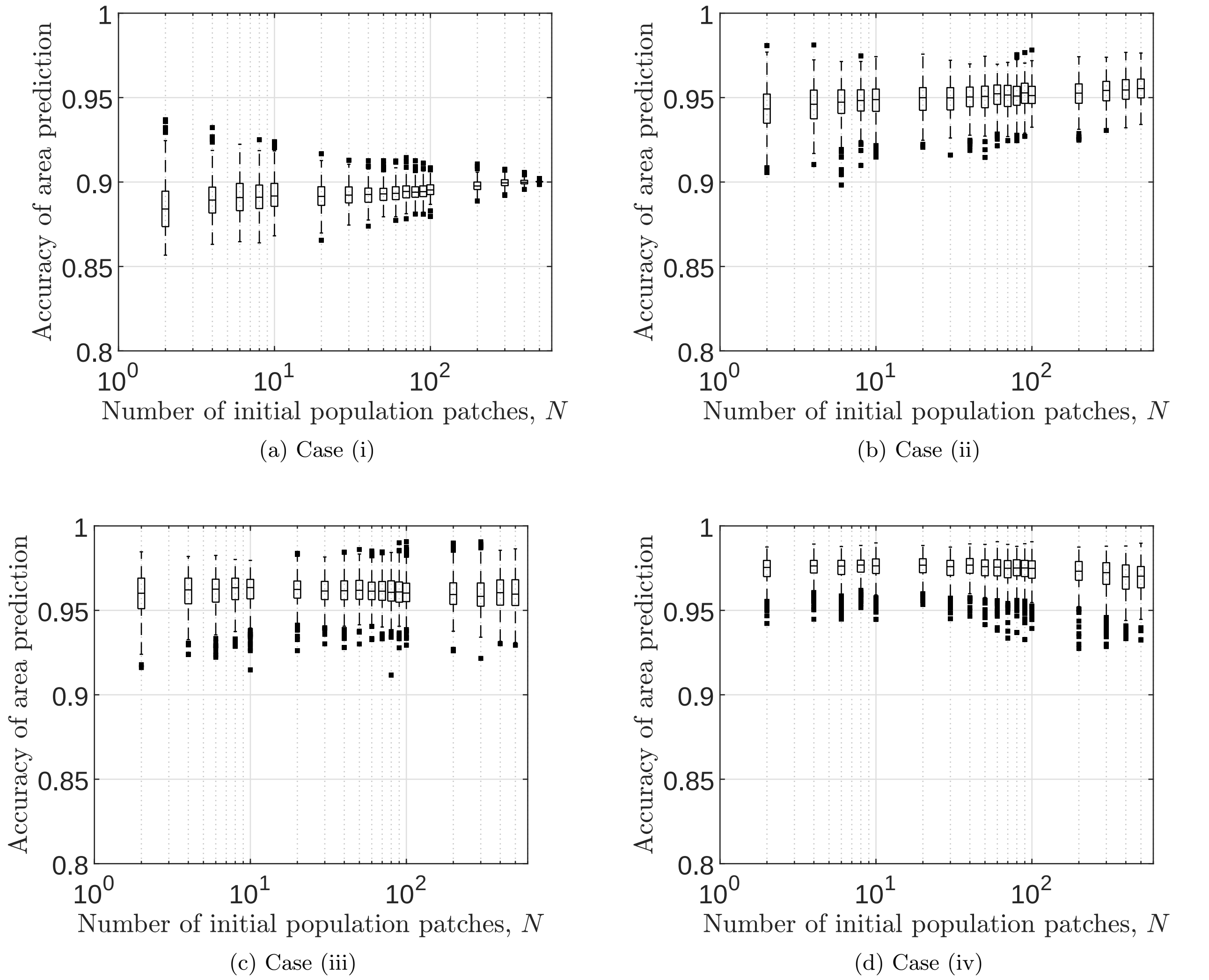


(a) Case (i)

(b) Case (ii)

(c) Case (iii)

(d) Case (iv)

Figure S3.1: **Accuracy of predictions of area covered for identical species.** The linear correlation between $A_{\text{pred}}$ and $A_{\text{sim}}$ is shown for each initial population density used in the Monte Carlo approach for each of the parameter landscape cases.

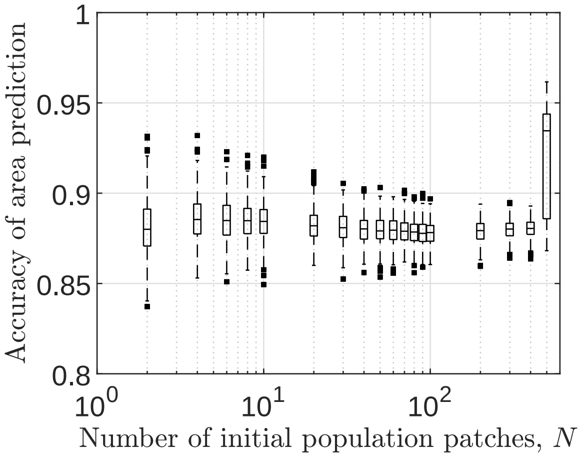


(a) Case (i)

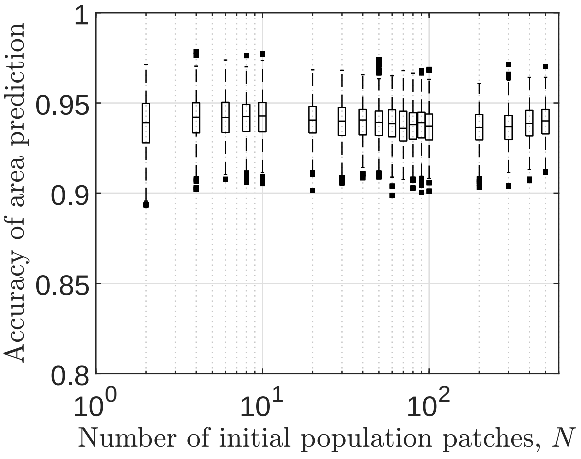


(b) Case (ii)

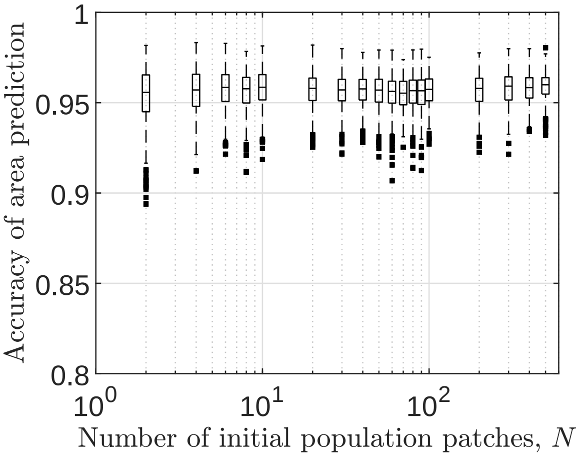


(c) Case (iii)

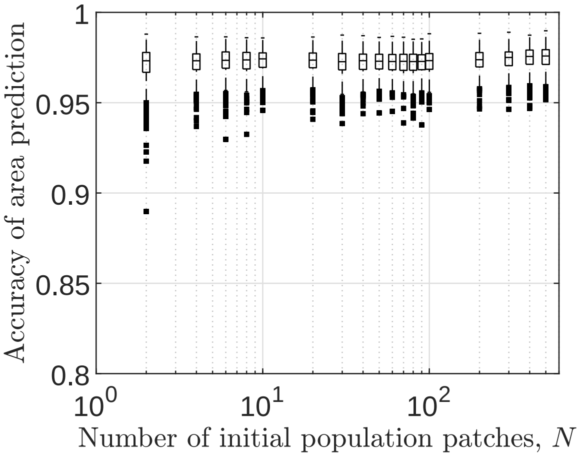


(d) Case (iv)

Figure S3.2: **Accuracy of predictions of area covered for antagonistic species.** The linear correlation between $A_{\text{pred}}$ and $A_{\text{sim}}$ is shown for each initial population density used in the Monte Carlo approach for each of the parameter landscape cases.

The region expected to be occupied by species $B_i$ during the finite time interval $[0, t]$ is

$$\Delta_{B_i}(t) = \left\{ \mathbf{x} \in \Delta_{B_i}^{\Omega} : \min_{\mathbf{y} \in \mathcal{B}_i} t_{\mathrm{FP}}^{(i)}(\mathbf{x}, \mathbf{y}) \leq t \right\}, \quad i = 1, \ldots, K.$$

Finally, the Voronoi index of species $B_i$, denoted by $V_i(t)$, is given by

$$V_i(t) := \frac{\mathrm{Area}\,(\Delta_{B_i}(t))}{\sum_{j=1}^{K} \mathrm{Area}\,\left(\Delta_{B_j}(t)\right)}, \quad i = 1, \ldots, K.$$

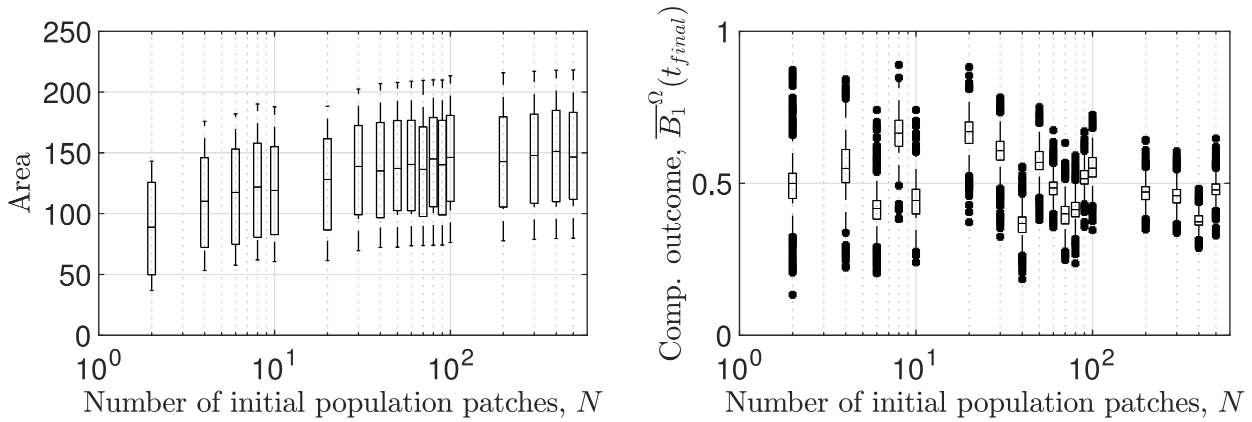


Figure S5.1: **Changes in spatial landscape induce variability in competitive outcome and area covered by range expansion if the initial population distribution is fixed**. The relations between initial population density and area covered by range expansion is shown in (a); the relation between initial population density and competitive outcome is shown in (b). Both cases show the full data of our Monte Carlo approach in which the initial population distribution is fixed for each initial population density and only the spatial landscape is randomly chosen for each model realisation.

### S5 Supplementary figures

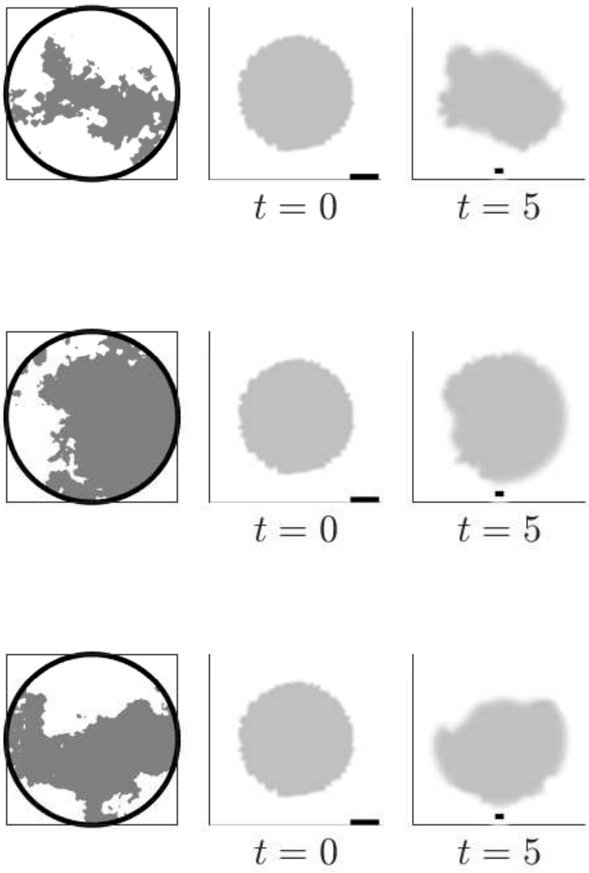


Figure S5.2: **Spatial homogeneity is preserved in range expansion of identical species.** Three independent model realisations of the theoretical framework with two identical species are shown each for high initial population density, represented by spatially uniform initial conditions. The left column of each subfigure shows the spatially heterogeneous parameter landscape; the middle column shows the initial condition of the system; the right column visualises the system at $t = t_{\text{final}} = 5$; the plot of the initial condition only shows a blow-up of the centre of the whole computational domain (see the black scale bars which are one unit length long). Grey colour corresponds to equal densities of $B_1$ and $B_2$.

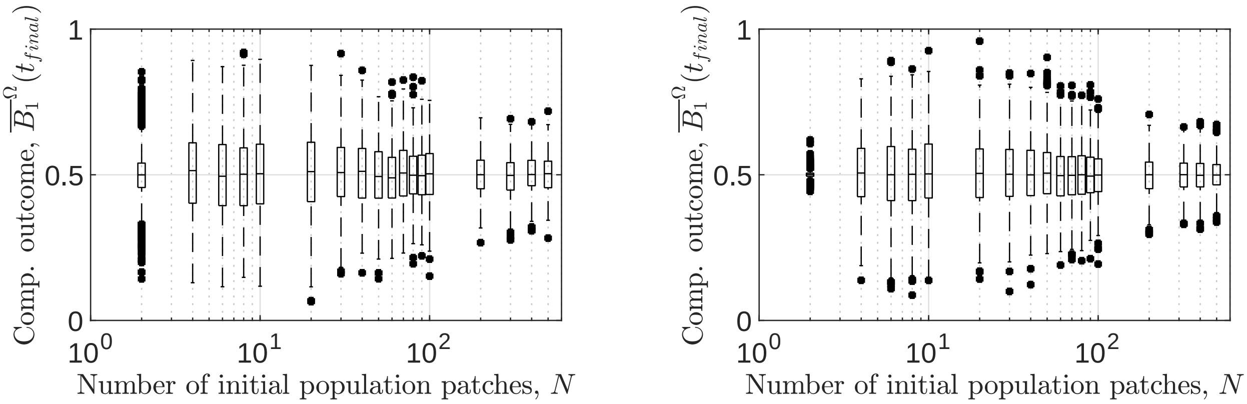


(a) Heterogeneous landscape.

(b) Homogeneous landscape.

Figure S5.3: **Initial population density is the main determinant of variability in competitive outcome for identical species.** The relation between initial population density and competitive outcome $\overline{B_i}^{\Omega}(t_{\text{final}})$ is shown for heterogeneous landscapes (a) and homogeneous landscapes (b) in the case of identical species. Part (b) is adapted from Eigentler et al. 2021.

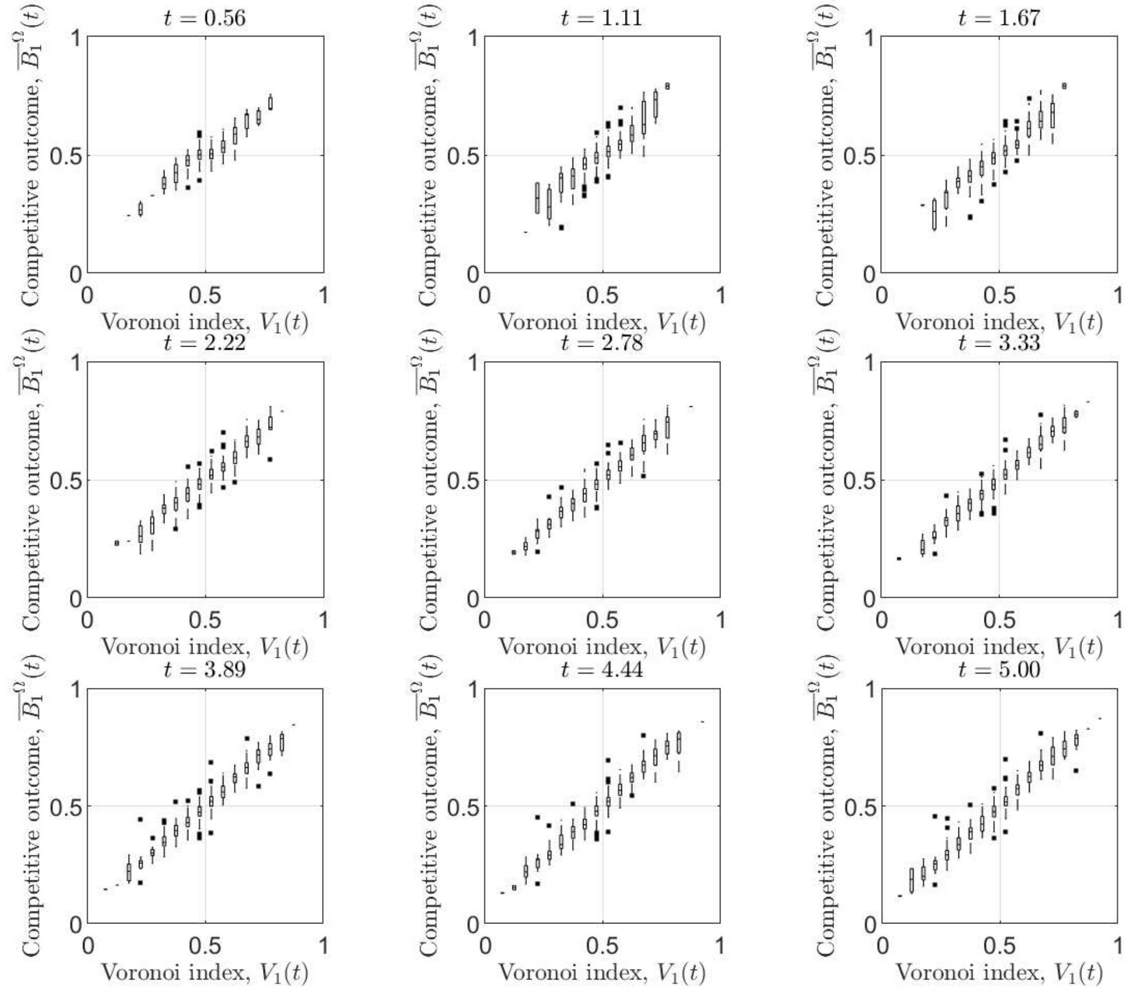


Figure S5.4: **Voronoi indices predict competitive outcome at all times.** The relation between the Voronoi index $V_1(t)$ and competitive outcome $\overline{B_1}^{\Omega}(t)$ is shown for a selection of times $t$.

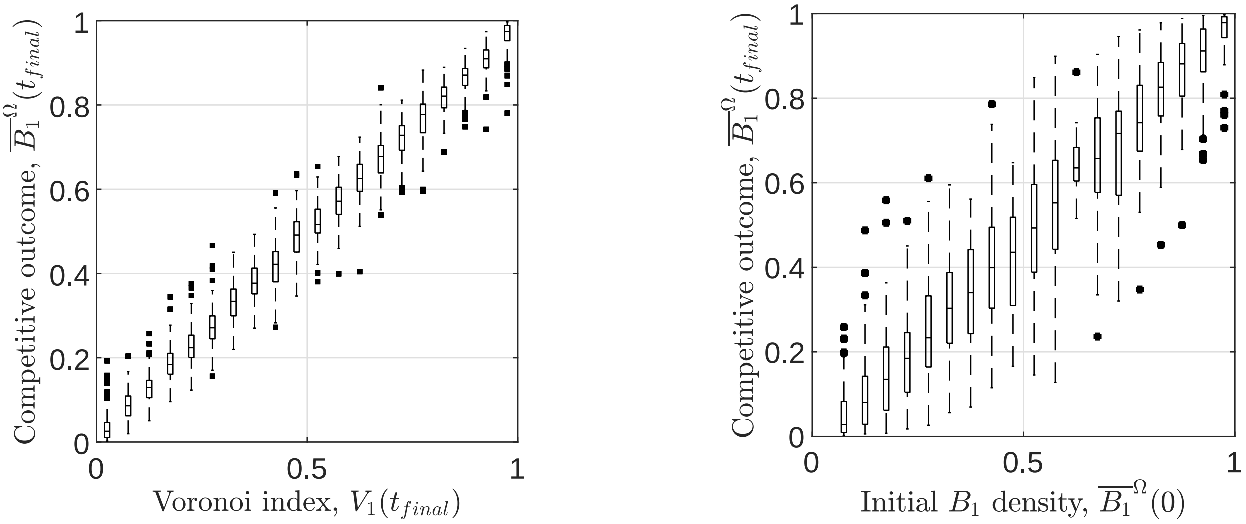


(a) Voronoi index determines competitive outcome.

(b) Competitive outcome is correlated with initial species ratio.

Figure S5.5: **Voronoi index determines competitive outcome independent of initial population ratio.** The relation between the Voronoi index and competitive outcome for randomly chosen initial population ratio (a) is compared with the relation between the initial population ratio and competitive outcome (b). The initial population density is fixed at $N = 20$ and the heterogeneous parameter landscape is created according to case (iii). Both the Voronoi index axis (a) and the initial population ratio axis (b) are binned for visualisation purposes but represent continuous variables.

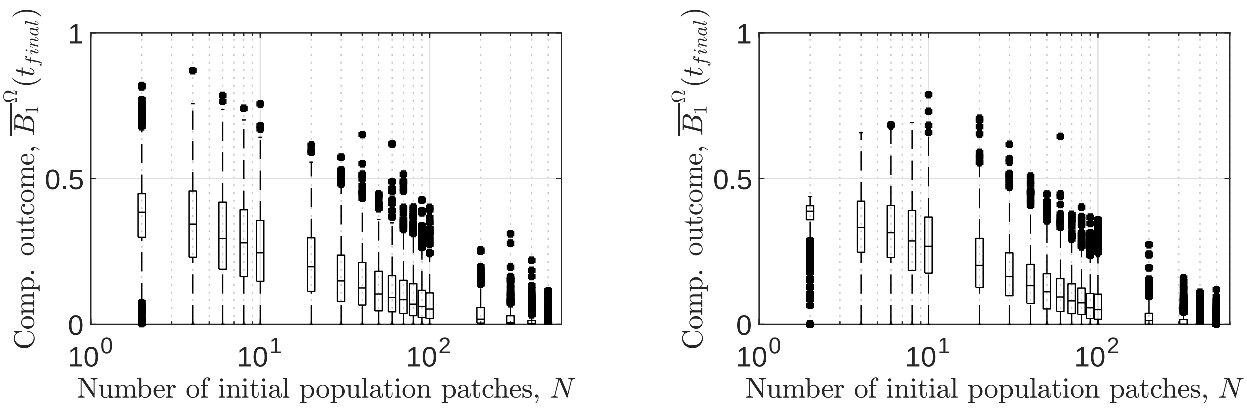


(a) Heterogeneous landscape.

(b) Homogeneous landscape.

Figure S5.6: **Initial population density is the main determinant of variability in competitive outcome for antagonistic species.** The relation between initial population density and competitive outcome $\overline{B_i}^{\Omega}(t_{\text{final}})$ is shown for heterogeneous landscapes (a) and homogeneous landscapes (b) in the case of antagonistic species. Part (b) is adapted from Eigentler et al. 2021.

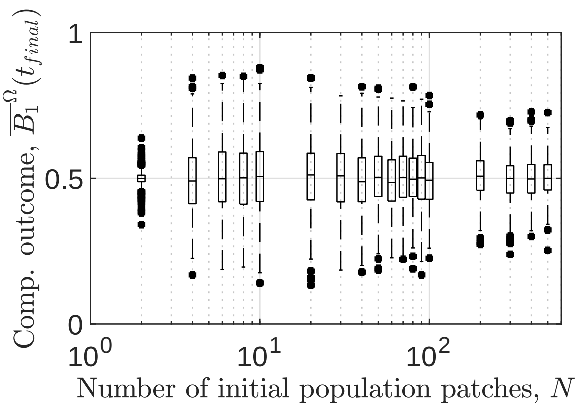


(a)

Figure S5.7: **Competitive outcome for two identical species as a function of initial population density in heterogeneous landscapes case (i).** For a full figure caption see the corresponding figure captions for case (iii) in the main text. The case-specific parameters are identical to those used for case (iii) in the main text.

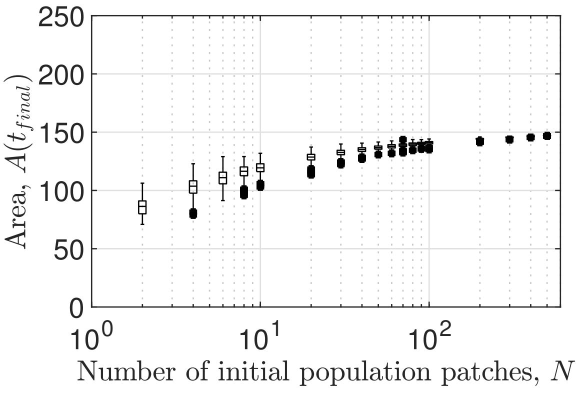


Figure S5.8: **Spatial heterogeneities determine variability in area covered by range expansion in heterogeneous landscapes case (i).** For a full figure caption see the corresponding figure captions for case (iii) in the main text. The case-specific parameters are identical to those used for case (iii) in the main text.

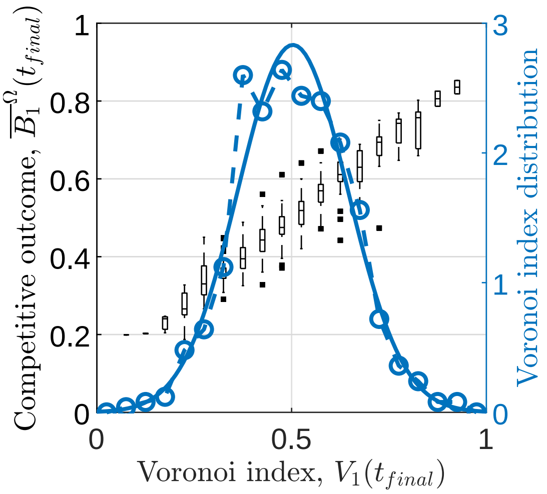


(a) Voronoi index as a predictor: $N = 6$

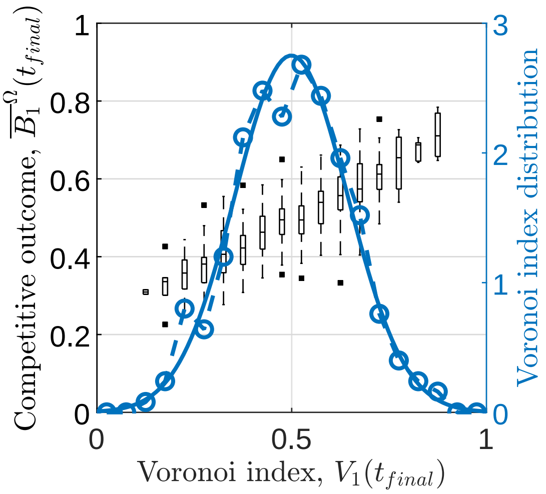


(b) Voronoi index as a predictor: $N = 100$

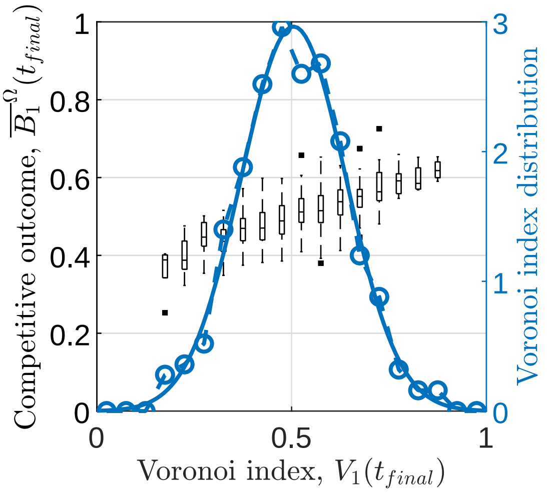


(c) Voronoi index as a predictor: $N = 500$

Figure S5.9: **Voronoi index as a predictor of competitive outcome for two identical species in heterogeneous landscapes case (i)**. For a full figure caption see the corresponding figure captions for case (iii) in the main text. The case-specific parameters are identical to those used for case (iii) in the main text.

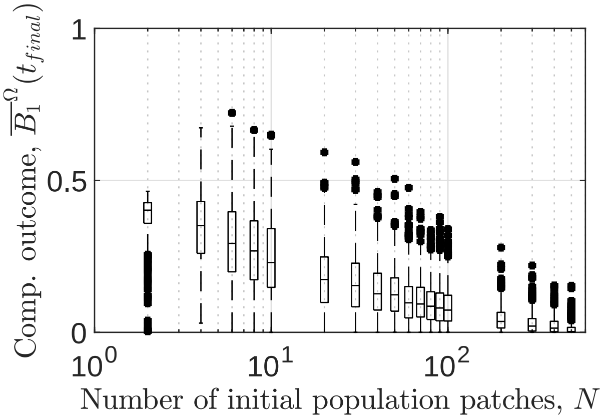


Figure S5.10: **Competitive outcome of two antagonistic species as a function of initial population density in heterogeneous landscapes case (i)**. For a full figure caption see the corresponding figure captions for case (iii) in the main text. The case-specific parameters are identical to those used for case (iii) in the main text.

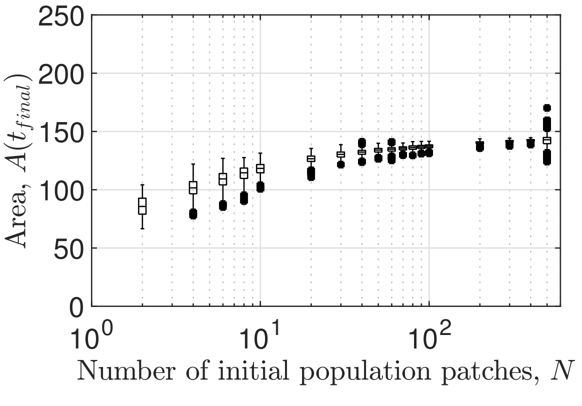


Figure S5.11: **Spatial heterogeneities determine variability in area covered by range expansion of antagonistic species in heterogeneous landscapes case (i)**. For a full figure caption see the corresponding figure captions for case (iii) in the main text.. The case-specific parameters are identical to those used for case (iii) in the main text.

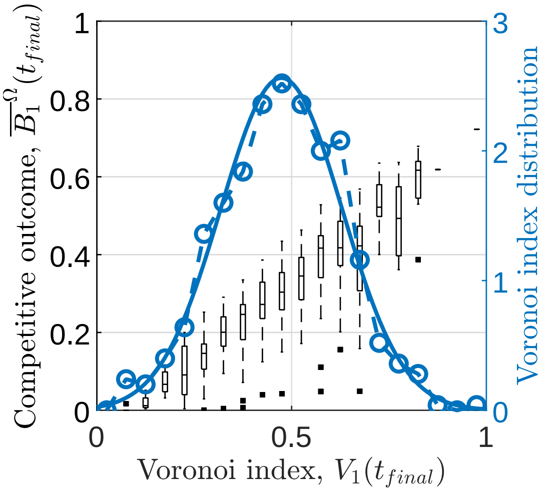


(a) Voronoi index as a predictor: $N = 6$

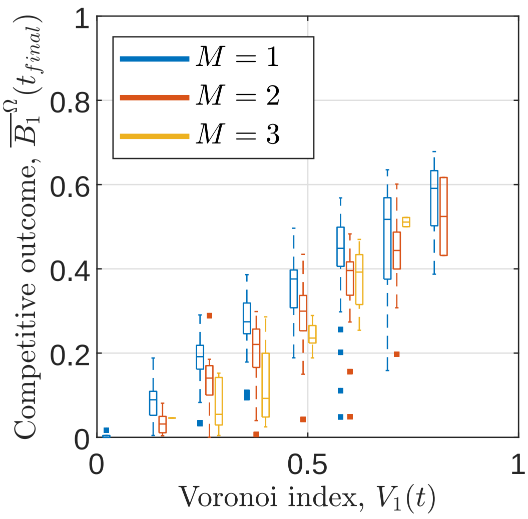


(b) Intraspecies connectedness as a filter for predictions: $N = 6$

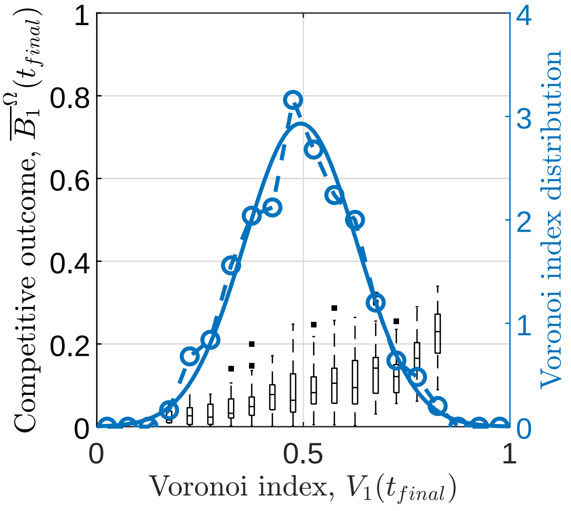


(c) Voronoi index as a predictor: $N = 100$

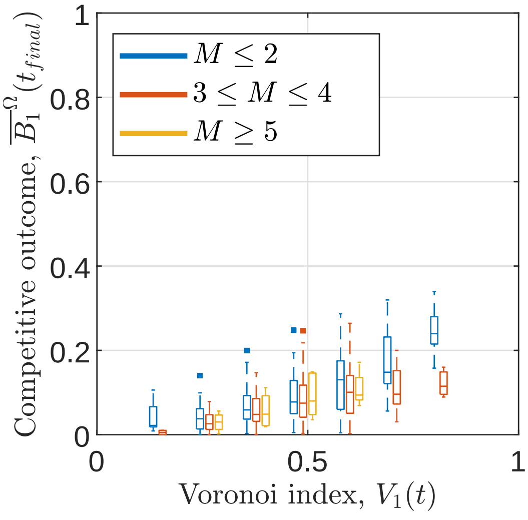


(d) Intraspecies connectedness as a filter for predictions: $N = 100$

Figure S5.12: **Voronoi index as a predictor of competitive outcome for two antagonistic species in heterogeneous landscapes case (i)**. For a full figure caption see the corresponding figure captions for case (iii) in the main text.. The case-specific parameters are identical to those used for case (iii) in the main text.

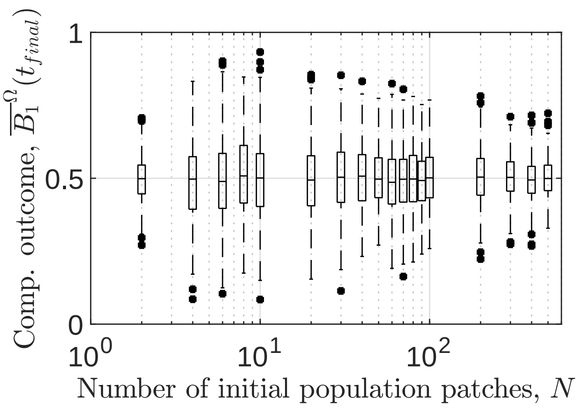


(a)

Figure S5.13: **Competitive outcome for two identical species as a function of initial population density in heterogeneous landscapes case (ii)**. For a full figure caption see the corresponding figure captions for case (iii) in the main text. The case-specific parameters are identical to those used for case (iii) in the main text.

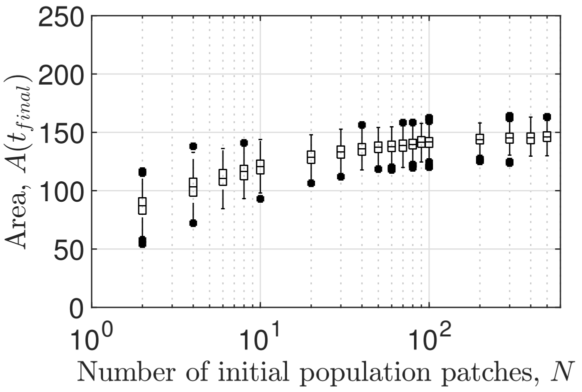


Figure S5.14: **Spatial heterogeneities determine variability in area covered by range expansion in heterogeneous landscapes case (ii)**. For a full figure caption see the corresponding figure captions for case (iii) in the main text. The case-specific parameters are identical to those used for case (iii) in the main text.

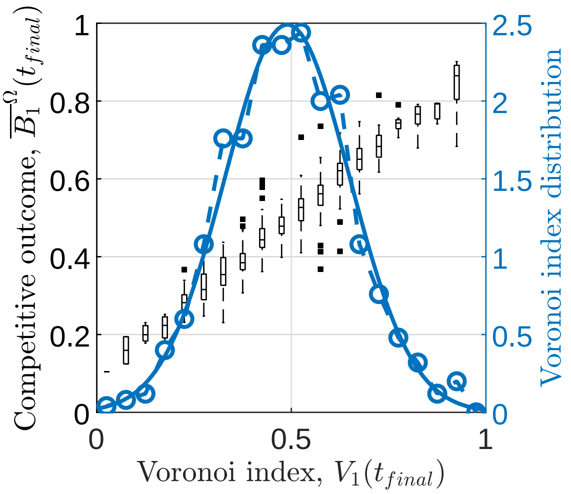


(a) Voronoi index as a predictor: $N = 6$

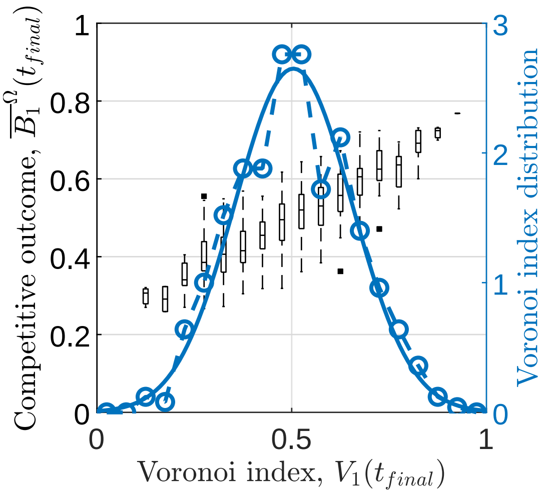


(b) Voronoi index as a predictor: $N = 100$

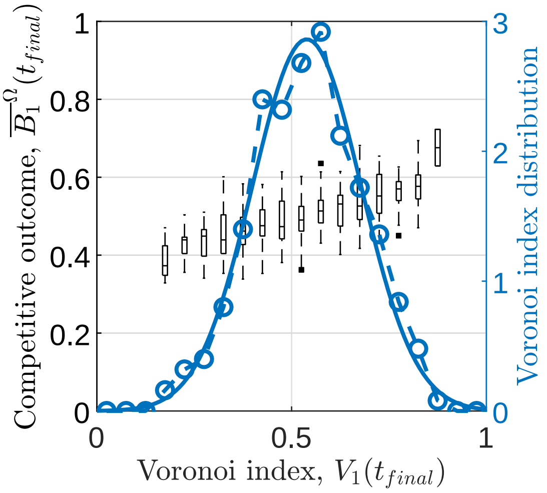


(c) Voronoi index as a predictor: $N = 500$

Figure S5.15: **Voronoi index as a predictor of competitive outcome for two identical species in heterogeneous landscapes case (ii)**. For a full figure caption see the corresponding figure captions for case (iii) in the main text. The case-specific parameters are identical to those used for case (iii) in the main text.

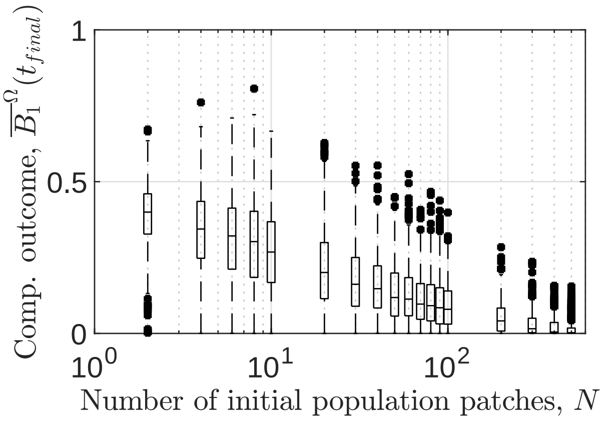


Figure S5.16: **Competitive outcome of two antagonistic species as a function of initial population density in heterogeneous landscapes case (ii)**. For a full figure caption see the corresponding figure captions for case (iii) in the main text. The case-specific parameters are identical to those used for case (iii) in the main text.

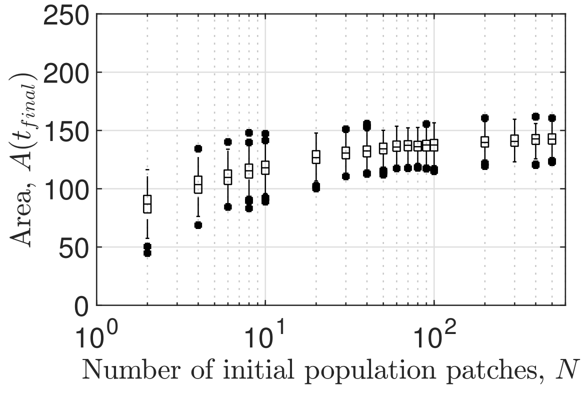


Figure S5.17: **Spatial heterogeneities determine variability in area covered by range expansion of antagonistic species in heterogeneous landscapes case (ii)**. For a full figure caption see the corresponding figure captions for case (iii) in the main text. The case-specific parameters are identical to those used for case (iii) in the main text.

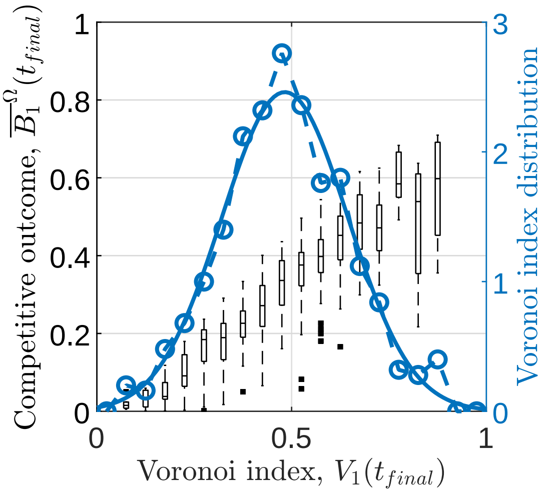


(a) Voronoi index as a predictor: $N = 6$

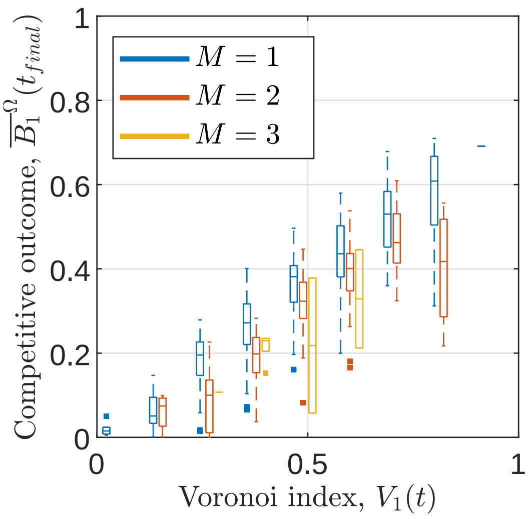


(b) Intraspecies connectedness as a filter for predictions: $N = 6$

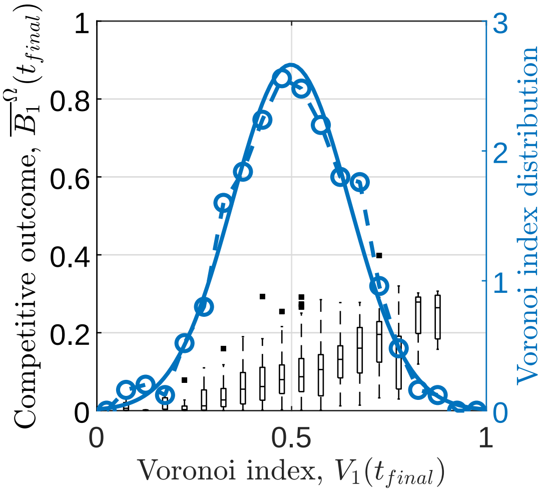


(c) Voronoi index as a predictor: $N = 100$

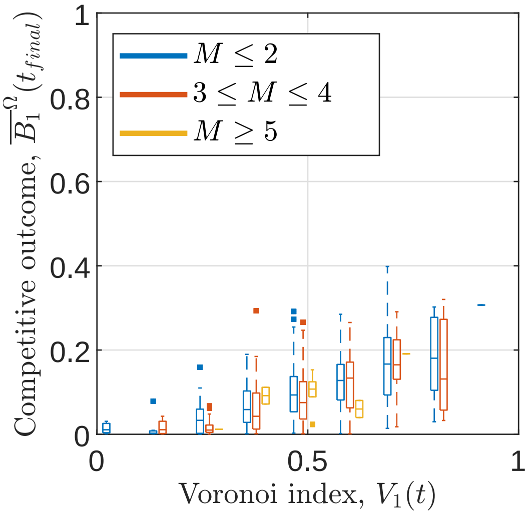


(d) Intraspecies connectedness as a filter for predictions: $N = 100$

Figure S5.18: **Voronoi index as a predictor of competitive outcome for two antagonistic species in heterogeneous landscapes case (ii)**. For a full figure caption see the corresponding figure captions for case (iii) in the main text. The case-specific parameters are identical to those used for case (iii) in the main text.

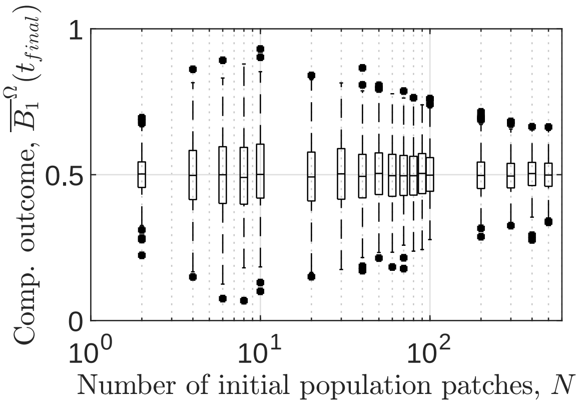


Figure S5.19: **Competitive outcome for two identical species as a function of initial population density in heterogeneous landscapes case (iv)**. For a full figure caption see the corresponding figure captions for case (iii) in the main text. The case-specific parameters to transform the monofractal $L$ into a heterogeneous landscape of model parameters are $r_i^{\text{mean}} = d_i^{\text{mean}} = 1$, $r^{\text{scale}} = 1$, $d^{\text{scale}} = 0.02$.

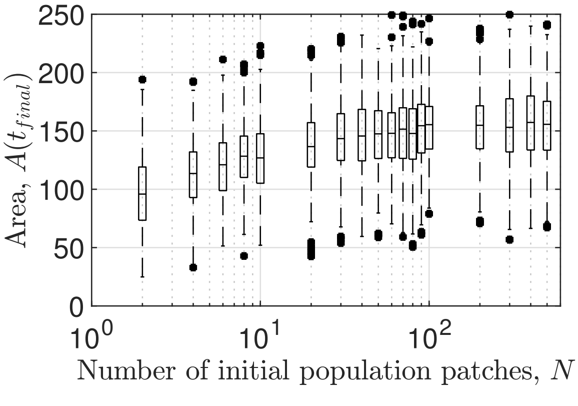


Figure S5.20: **Spatial heterogeneities determine variability in area covered by range expansion in heterogeneous landscapes case (iv)**. For a full figure caption see the corresponding figure captions for case (iii) in the main text. The case-specific parameters to transform the monofractal $L$ into a heterogeneous landscape of model parameters are $r_i^{\text{mean}} = d_i^{\text{mean}} = 1$, $r^{\text{scale}} = 1$, $d^{\text{scale}} = 0.02$.

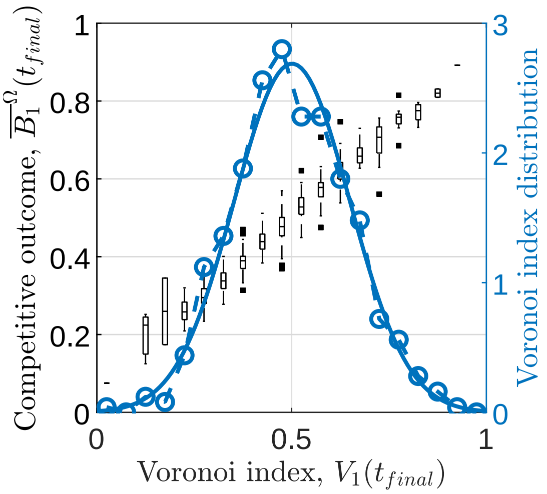


(a) Voronoi index as a predictor: $N = 6$

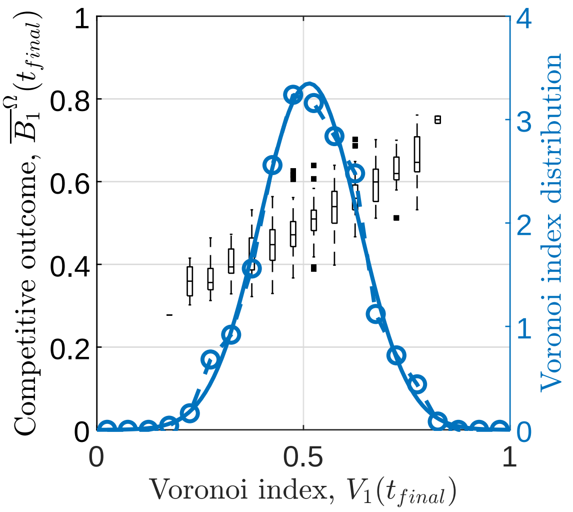


(b) Voronoi index as a predictor: $N = 100$

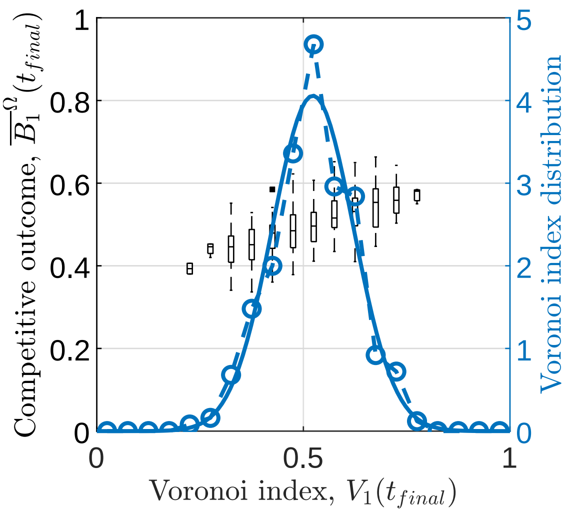


(c) Voronoi index as a predictor: $N = 500$

Figure S5.21: **Voronoi index as a predictor of competitive outcome for two identical species in heterogeneous landscapes case (iv)**. For a full figure caption see the corresponding figure captions for case (iii) in the main text. The case-specific parameters to transform the monofractal $L$ into a heterogeneous landscape of model parameters are $r_i^{\text{mean}} = d_i^{\text{mean}} = 1$, $r^{\text{scale}} = 1$, $d^{\text{scale}} = 0.02$.

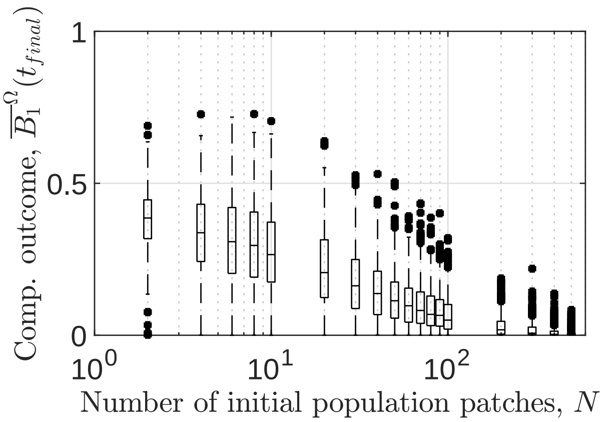


Figure S5.22: **Competitive outcome of two antagonistic species as a function of initial population density in heterogeneous landscapes case (iv)**. For a full figure caption see the corresponding figure captions for case (iii) in the main text. The case-specific parameters to transform the monofractal $L$ into a heterogeneous landscape of model parameters are $r_i^{\text{mean}} = d_i^{\text{mean}} = 1$, $r^{\text{scale}} = 1$, $d^{\text{scale}} = 0.02$.

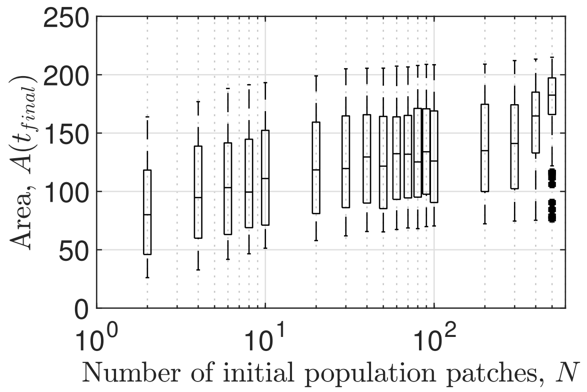


Figure S5.23: **Spatial heterogeneities determine variability in area covered by range expansion of antagonistic species in heterogeneous landscapes case (iv)**. For a full figure caption see the corresponding figure captions for case (iii) in the main text. The case-specific parameters to transform the monofractal $L$ into a heterogeneous landscape of model parameters are $r_i^{\text{mean}} = d_i^{\text{mean}} = 1$, $r^{\text{scale}} = 1$, $d^{\text{scale}} = 0.02$.

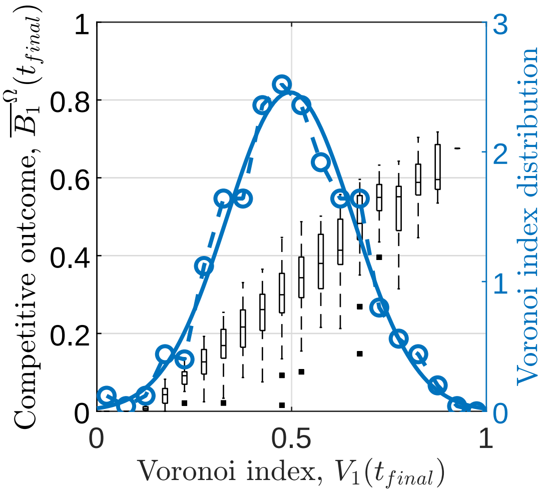


(a) Voronoi index as a predictor: $N = 6$

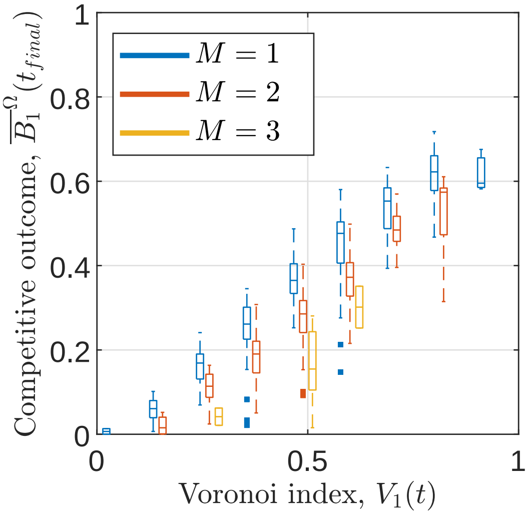


(b) Intraspecies connectedness as a filter for predictions: $N = 6$

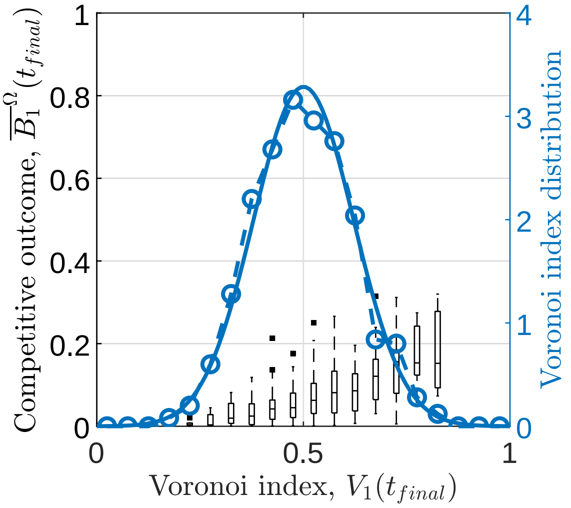


(c) Voronoi index as a predictor: $N = 100$

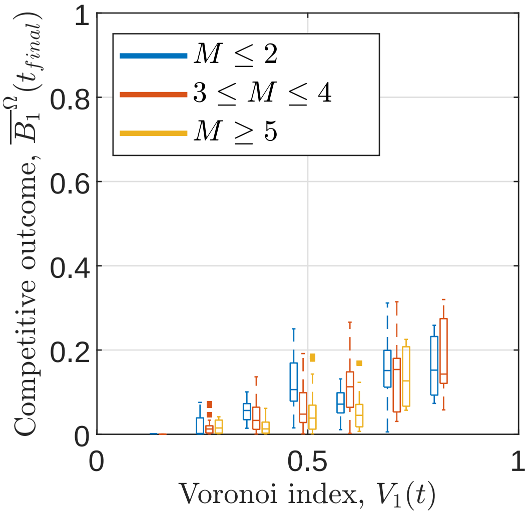


(d) Intraspecies connectedness as a filter for predictions: $N = 100$

Figure S5.24: **Voronoi index as a predictor of competitive outcome for two antagonistic species in heterogeneous landscapes case (iv)**. For a full figure caption see the corresponding figure captions for case (iii) in the main text. The case-specific parameters to transform the monofractal $L$ into a heterogeneous landscape of model parameters are $r_i^{\text{mean}} = d_i^{\text{mean}} = 1$, $r^{\text{scale}} = 1$, $d^{\text{scale}} = 0.02$.
